# A history of avoidance does not impact extinction learning in male rats

**DOI:** 10.1101/2023.09.22.558816

**Authors:** Alba López-Moraga, Laura Luyten, Tom Beckers

## Abstract

Pervasive avoidance is one of the central symptoms of all anxiety-related disorders. In treatment, avoidance behaviors are typically discouraged because they are assumed to maintain anxiety. Yet, it is not clear that engaging in avoidance is always detrimental. In this study, we used a platform-mediated avoidance task to investigate the influence of avoidance history on extinction learning in male rats. Our results show that having the opportunity to avoid during fear acquisition training has no marked effect on the extinction of auditory cued fear in a platform-mediated avoidance procedure that constitutes a realistic approach/avoidance conflict in male rats, regardless of whether avoidance was possible during extinction or not. This suggests that imposing a realistic cost on avoidance behavior prevents the adverse effects that avoidance has been claimed to have on extinction, but even then, avoidance does not appear to have clear positive effects on extinction learning nor on retention either.

## Introduction

Pervasive avoidance is one of the central symptoms of all anxiety-related disorders^1^. Avoidance behavior can be broadly defined as any external or internal response that increases the distance between an individual and a (perceived or actual) threat or aversive event. As such, avoidance behaviors can take subtle forms (e.g., a person with social anxiety disorder may rehearse what to say or may speak quietly)^2^. In clinical management, individuals with an anxiety-related disorder are typically discouraged from engaging in avoidance because it is assumed to maintain anxiety^3^. In line with this notion, in laboratory studies in humans, conditioned fear that is established by pairing a neutral cue with an aversive or threatening outcome will reduce when that neutral cue is repeatedly presented without the threatening outcome (i.e., extinction training), but when individuals are given the opportunity to perform avoidance behaviors during extinction training, such extinction training will be less effective^4^. Correspondingly, rodent research has reported that instrumental avoidance can result in lesser extinction^5^.

Yet, despite the widely accepted notion that avoidance can maintain anxiety, it is not clear that avoidance is always detrimental^6^. It has been reported that the ability to rely on safety behaviors can increase the willingness of patients to endure exposure treatment^7^, which may be important given that the anxiety experienced during exposure-based therapy is one of the reasons that patients end treatment prematurely or never even start such streatment^2^. For example, dropout from prolonged exposure therapy was examined in a sample of 2606 patients. Three out of ten patients completed less than 8 sessions, which is considered a minimum therapeutic dose for most patients. Clinicians attributed dropout to distress or avoidance in 45% of the cases^8^. Moreover, research has suggested that engaging in safety behaviors does not interfere with treatment outcome^9^ and that it could provide a sense of control that can even yield more effective exposure treatment^10^. Supporting this premise, prior studies have found that a history of controllability over stressors may enhance extinction of conditioned fear in humans^11^ and rats^12^.

In sum, while sustained avoidance throughout extinction training may be largely detrimental for fear reduction, having the opportunity to avoid prior to or in the initial stages of extinction training may actually improve subsequent fear reduction. It is that prediction that we set out to test here using a platform-mediated avoidance (PMA) task. In this task, food-restricted rats are first trained to press a lever for food reward. When fear acquisition starts, rats learn that they can avoid a tone-signaled foot shock by stepping onto a platform during the tone, at the expense of lever-pressing for food^13–15^. Thus, fearful avoidance competes with food-seeking behavior, which mirrors the clinical reality of anxious individuals who avoid threatening but also possibly rewarding situations, constituting an approach-avoidance conflict^16^.This translational characteristic differentiates PMA from other active avoidance procedures (e.g., shuttle avoidance) that have no cost associated with the avoidance response^17^. We evaluated different hypotheses regarding the possible influence of a history of avoidance on extinction, exploiting the flexibility that characterizes this task.

Based on prior work suggesting that a sense of control could promote better extinction^11,12^, we hypothesized that a history of successful avoidance might promote the later learning of safety during extinction training^18^. In Experiment 1, we hypothesized that rats that had had a platform present during fear acquisition would learn extinction faster than their Yoked counterparts. For Experiment 2, the task was adapted so that rats with a history of avoidance had the possibility to perform avoidance responses also during extinction training. Our objective was to investigate whether rats would exhibit sustained avoidance despite the omission of foot shocks. Previous animal studies have shown that avoidance tends to persist following extinction training^19^, especially when avoidance was also possible during extinction^5^, owing to the fact that avoidance responses during extinction training reduce the possibility for the animal to experience that CSs are no longer followed by a US in the extinction phase. Finally, in Experiment 3, we investigated the longevity of the effect of avoidance on extinction by assessing spontaneous recovery in Avoiders and Yoked controls.

## Results

### Two days of training are sufficient for avoidance learning

After learning to lever press on a VI 30 s reinforcement schedule, rats were divided into two groups: Avoiders (A) and Yoked controls (Y). In all three experiments, rats in group A could avoid the US that co-terminated with the CS by stepping onto a platform prior to US onset, whereas rats in group Y were presented with a CS-US contingency that was determined by the actions of their companion animal in group A. As such, animals in both groups experienced the exact same CS-US contingencies.

As expected, two days of avoidance training were sufficient for the rats in group A to learn to avoid a foot shock by stepping onto the platform in response to the CS tone (Fig 1) (Experiment 1: V = 1, p<0.001; Experiment 2: V = 1, p<0.001, Experiment 3: t(11) = −2.36, p = 0.04, see Table 1 for all statistical tests), also when avoidance training was preceded by purely Pavlovian training in Experiment 3 (Fig S1 and Table S1). Furthermore, in both groups, we found a significant increase in freezing from day 1 to day 2 in Experiments 1 (F(1, 22) = 59.80, p < 0.001) and 2 (Q(1, 10.70) = 23.33, p < 0.001). Such an increase was not observed in Experiment 3, where we did find significantly higher freezing in the Avoider rats compared to their Yoked counterparts (F(1, 21) = 5.85, p = 0.025). Results for suppression of lever pressing and rearing can be found in Fig S2 and Table S2.

**Fig 1.**
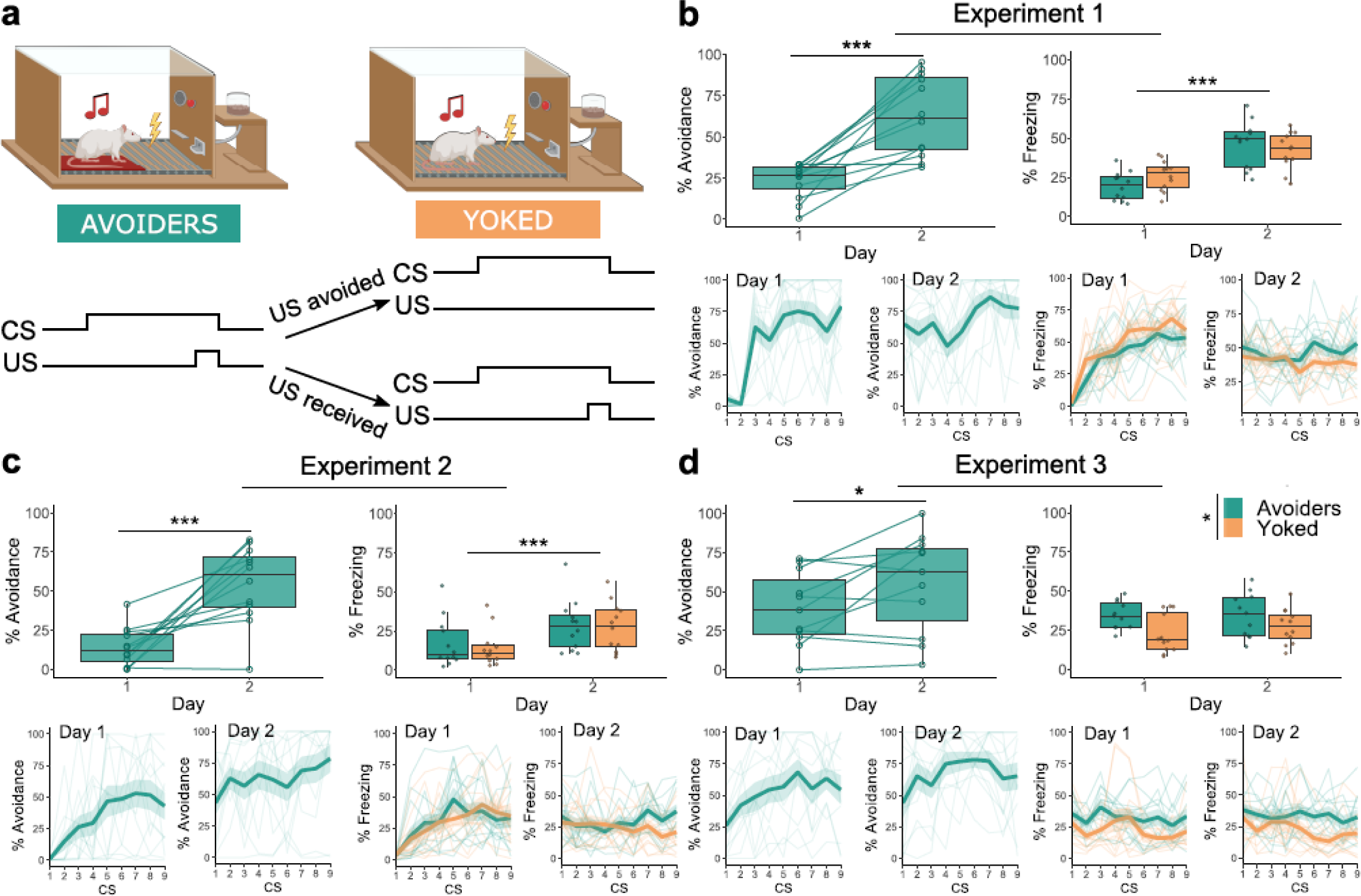
Avoidance training results for Experiments 1-3. The box plots represent the average of the first 3 CSs on each day. The bold lines in the trial-by-trial plots represent the mean and the surrounding shaded area the standard error of the mean. Results are expressed in % of time during CS presentations. **a.** Graphic representation of the avoidance training sessions in the three experiments. **b.** In the Avoider group, avoidance increased significantly from day 1 to 2 (V = 1, p < 0.001, r = 0.861). Across both groups, freezing increased significantly from day 1 to 2 (F(1, 22) = 59.80, p < 0.001, η_p_^2^ = 0.731). **c.** In the Avoider group, avoidance increased significantly from day 1 to 2 (V = 1, p < 0.001, r = 0.861). Across both groups, freezing increased significantly from day 1 to 2 (Q(1, 10.70) = 23.33, p < 0.001, η_p_^2^ = 0.477). **d.** In the Avoider group, avoidance increased significantly from day 1 to 2 (t(11) = −2.74, p = 0.019, d = −0.791). Freezing differed significantly between Avoiders and Yoked animals (F(1, 22) = 6.66, p = 0.017, η_p_^2^ = 0.232). *p□<□0.05, ***p□<□0.001.

**Table 1.** Avoidance training.

| Experiment | Measure | Statistical Test | Result | Effect size |
| --- | --- | --- | --- | --- |
| 1 | Avoidance | Wilcoxon signed ranks test | Day: $V=1$, **$p < 0.001$** | $r = 0.861$ |
| | Freezing | Mixed ANOVA | Group: $F(1, 22) = 0.15$, $p = 0.701$<br>Day: $F(1, 22) = 59.80$, **$p < 0.001$**<br>G*D: $F(1, 22) = 3.63$, $p = 0.07$ | Group: $\eta_p^2 = 0.007$<br>Day: $\eta_p^2 = 0.731$<br>G*D: $\eta_p^2 = 0.142$ |
| 2 | Avoidance | Wilcoxon signed ranks test | Day: $V=1$, **$p < 0.001$** | $r = 0.861$ |
| | Freezing | Non-parametric mixed ANOVA | Group: $Q(1, 12.77) = 0.04$, $p = 0.84$<br>Day: $Q(1, 10.70) = 23.33$, **$p < 0.001$**<br>G*D: $Q(1, 10.70) = 0.24$, $p = 0.63$ | Group: $\eta_p^2 = 0.005$<br>Day: $\eta_p^2 = 0.477$<br>G*D: $\eta_p^2 = 0.008$ |
| 3 | Avoidance | Paired t-test | Day: $t(11) = -2.36$, **$p = 0.04$** | $d = -0.711$ |
| | Freezing | Mixed ANOVA | Group: $F(1, 21) = 5.85$, **$p = 0.025$**<br>Day: $F(1, 21) = 0.58$, $p = 0.453$<br>G*D: $F(1, 21) = 0.62$, $p = 0.441$ | Group: $\eta_p^2 = 0.218$<br>Day: $\eta_p^2 = 0.027$<br>G*D: $\eta_p^2 = 0.029$ |

In sum, as hypothesized, rats in group A readily acquired signaled avoidance in just two days of training. In addition, we observed that rats in both groups developed fear-related behaviors such as CS-elicited freezing and suppression of lever pressing over the training days.

### A history of avoidance does not impact the capacity for extinction learning

After avoidance learning, extinction sessions were performed in all three experiments. Experiment 1 and 2 involved extinction sessions on 4 consecutive days. In Experiment 1 the platform was absent, whereas in Experiment 2 it was present for both groups. In Experiment 3, extinction training lasted for 2 days only, with the platform present. Despite the differences in design, the three experiments yielded comparable results across the various dependent variables. In all cases, groups A and Y similarly showed extinction of fear responding to the CS (Fig 2).

**Fig 2.**
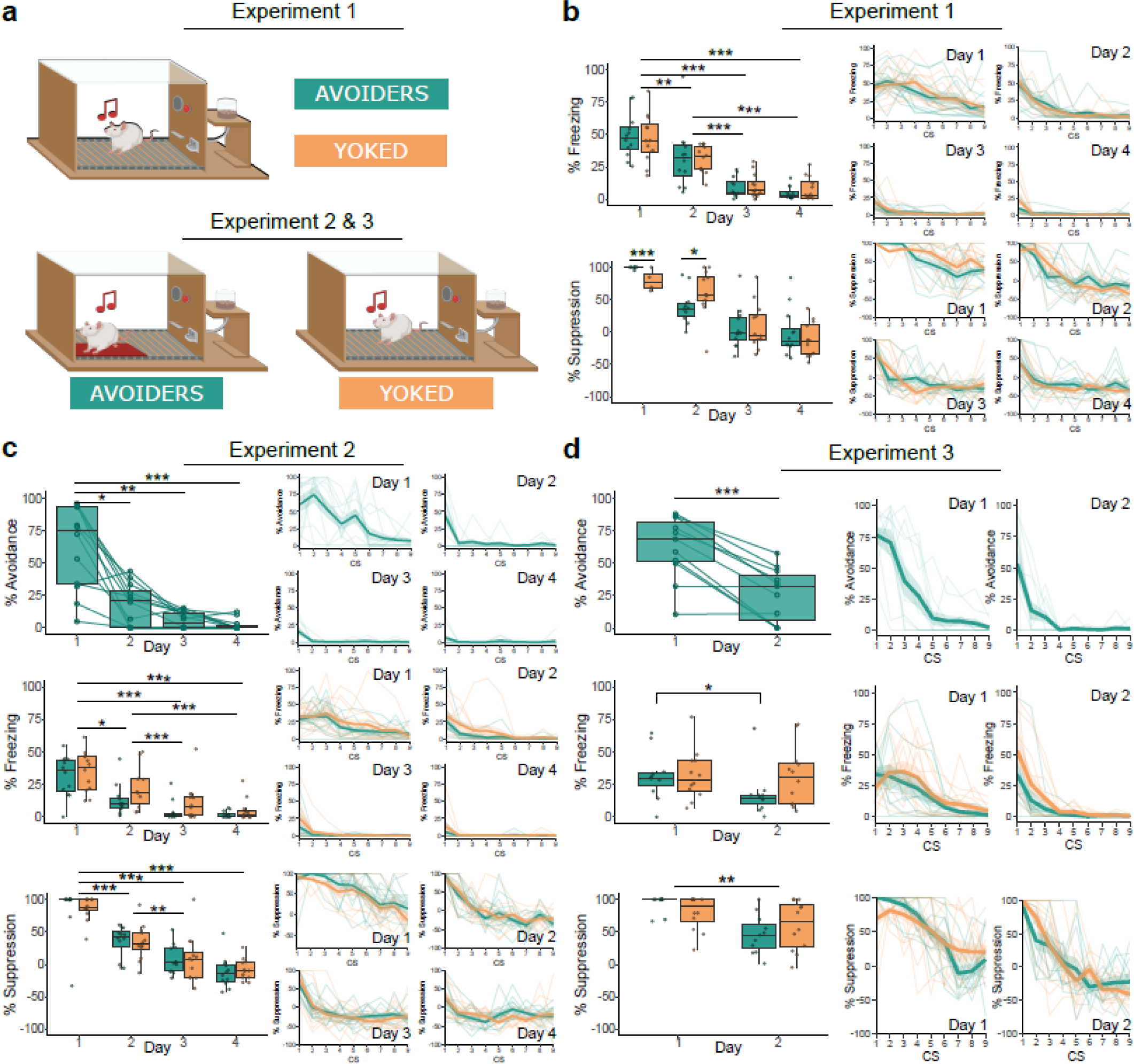
Extinction results for Experiments 1-3. The box plots represent the average of the first 3 CSs on each day. The bold lines in the trial-by-trial plots represent the mean and the surrounding shaded area the standard error of the mean. Results are expressed in % of time during CS presentations. **a.** Graphic representation of the extinction training sessions. **b.** Across both groups, freezing decreased significantly over the extinction sessions (Q(3, 11.33) = 87.44, p < 0.001, η_p_^2^ = 0.764). A further investigation of the interaction between group and extinction session (Q(3, 10.61) = 7.14, p = 0.007, η_p_^2^ = 0.13) revealed that on day 1 the Avoider group suppressed lever pressing more than the Yoked group (V= 135, p <0.001), whereas on day 2 the Yoked group shower more suppression of lever pressing than the Avoider group (V= 32.5, p = 0.024). **c.** In the Avoider group, avoidance decreased significantly over the extinction sessions (^2^(3) = 28, p<0.001, W = 0.779). Freezing decreased significantly across both groups (Q(3, 9.6) = 30.56, p < 0.001, η_p_^2^ = 0.694), as did lever pressing (Q(3, 10.98) = 258.58, p < 0.001, η_p_^2^ = 0.77). **d.** In the Avoider group, avoidance decreased significantly from day 1 to 2 (t(10) = 5.9, p < 0.001, d = 1.78). An interaction between group and test day (Q(1, 11.76) = 8.55, p = 0.013) revealed that freezing was significantly decreased in the Avoider group (V= 52, p = 0.014, r = 0.476), but not the Yoked group. *p□<□0.05, ** p < 0.01, ***p□<□0.001.

Avoidance could not be assessed in Experiment 1 (the platform was absent during extinction training), but in the other two experiments, animals in group A exhibited a significant reduction of avoidance after just one extinction session (Experiment 2: χ^2^(3) = 28, p < 0.001, Experiment 3: (t(10) = 5.9, p < 0.001, see Table 2 for all statistics).

**Table 2.** Extinction.

| Experiment | Measure | Statistical Test | Result | Effect size |
| --- | --- | --- | --- | --- |
| 1 | Freezing | Non-parametric mixed ANOVA | Group: $Q(1, 13.97) = 0.09$, $p = 0.765$<br>Day: $Q(3, 11.33) = 87.44$, **$p < 0.001$**<br>G*D: $Q(3, 11.33) = 0.19$, $p = 0.902$ | Group: $\eta_p^2 < 0.001$<br>Day: $\eta_p^2 = 0.764$<br>G*D: $\eta_p^2 = 0.011$ |
| | Freezing | Pairwise comparisons with Wilcoxon rank sum test, Bonferroni correction | Day 1-2: **$p = 0.005$**<br>Day 1-3: **$p < 0.001$**<br>Day 2-3: **$p < 0.001$**<br>Day 1-4: **$p < 0.001$**<br>Day 2-4: **$p < 0.001$**<br>Day 3-4: $p = 0.514$ | |
| | Suppression of lever pressing | Non-parametric mixed ANOVA | Group: $Q(1, 13.30) = 0.08$, $p = 0.775$<br>Day: $Q(3, 10.61) = 135.78$, **$p < 0.001$**<br>G*D: $Q(3, 10.61) = 7.14$, **$p = 0.007$** | Group: $\eta_p^2 = 0.001$<br>Day: $\eta_p^2 = 0.781$<br>G*D: $\eta_p^2 = 0.13$ |
| | Suppression of lever pressing | Simple effects by day: Wilcoxon signed ranks test | Day 1 – Group: $V= 135$, **$p < 0.001$**<br>Day 2 – Group: $V= 32.5$, **$p = 0.024$** | Day 1 – Group: $r = 0.77$<br>Day 2 – Group: $r = 0.466$ |
| | | | Day 3 – Group: $V = 74$, $p = 0.932$<br>Day 4 – Group: $V = 81$, $p = 0.63$ | Day 3 – Group: $r = 0.024$<br>Day 4 – Group: $r = 0.106$ |
| 2 | Avoidance | Friedman rank sum test | Day: $\chi^2(3) = 28$, $p < 0.001$ | $W = 0.779$ |
| | Avoidance | Pairwise comparisons with Wilcoxon rank sum test with Bonferroni correction | Day 1-2: $p = 0.017$<br>Day 1-3: $p = 0.001$<br>Day 2-3: $p = 0.714$<br>Day 1-4: $p < 0.001$<br>Day 2-4: $p = 0.124$<br>Day 3-4: $p = 0.612$ | |
| | Freezing | Non-parametric mixed ANOVA | Group: $Q(1, 13.4) = 2.90$, $p = 0.112$<br>Day: $Q(3, 9.6) = 30.56$, $p < 0.001$<br>G*D: $Q(3, 9.6) = 2.20$, $p = 0.153$ | Group: $\eta_p^2 = 0.087$<br>Day: $\eta_p^2 = 0.694$<br>G*D: $\eta_p^2 = 0.031$ |
| | Freezing | Pairwise comparisons with Wilcoxon rank sum test with Bonferroni correction | Day 1-2: $p = 0.011$<br>Day 1-3: $p < 0.001$<br>Day 2-3: $p = 0.001$<br>Day 1-4: $p < 0.001$<br>Day 2-4: $p < 0.001$<br>Day 3-4: $p = 0.514$ | |
| | Suppression of lever pressing | Non-parametric mixed ANOVA | Group: $Q(1, 13.16) = 0.41$, $p = 0.533$<br>Day: $Q(3, 10.98) = 258.58$, $p < 0.001$<br>G*D: $Q(3, 10.98) = 0.94$, $p = 0.455$ | Group: $\eta_p^2 < 0.001$<br>Day: $\eta_p^2 = 0.77$<br>G*D: $\eta_p^2 = 0.003$ |
| | Suppression of lever pressing | Pairwise comparisons with Wilcoxon rank sum test with Bonferroni correction | Day 1-2: $p < 0.001$<br>Day 1-3: $p < 0.001$<br>Day 2-3: $p = 0.004$<br>Day 1-4: $p < 0.001$<br>Day 2-4: $p < 0.001$<br>Day 3-4: $p = 0.123$ | |
| 3 | Avoidance | Paired t-test | Day: $t(10) = 5.9$, $p < 0.001$ | $d = 1.78$ |
| | Freezing | Non-parametric mixed ANOVA | Group: $Q(1, 7.7) = 1.36$, $p = 0.278$<br>Day: $Q(1, 11.76) = 17.46$, $p = 0.001$<br>G*D: $Q(1, 11.76) = 8.55$, $p = 0.013$ | Group: $\eta_p^2 = 0.037$<br>Day: $\eta_p^2 = 0.286$<br>G*D: $\eta_p^2 = 0.235$ |
| | Freezing | Simple effects by group: Wilcoxon signed ranks test | Avoiders – Day: $V = 52$, $p = 0.014$<br>Yoked – Day: $V = 47$, $p = 0.569$ | $r = 0.476$<br>$r = 0.082$ |
| | Suppression of lever pressing | Non-parametric mixed ANOVA | Group: $Q(1, 9.69) = 0.07$, $p = 0.796$<br>Day: $Q(1, 9.45) = 14.77$, $p = 0.004$ | Group: $\eta_p^2 < 0.001$<br>Day: $\eta_p^2 = 0.514$ |
| | | | G*D: $Q(1, 9.45) = 2.85, p = 0.124$ | G*D: $\eta_p^2 = 0.157$ |

Freezing was significantly reduced after one extinction session in Experiment 1 (Q(3, 11.33) = 87.44, p < 0.001) and Experiment 2 (Q(3, 9.6) = 30.56, p < 0.001); no significant differences between groups were found. In Experiment 3, there was a significant interaction effect (Q(1, 11.76) = 8.55, p = 0.013), and the analysis of simple main effects of day per group showed that animals in group A showed a significant reduction of freezing (V = 52, p = 0.014), whereas animals in group Y did not (V = 47, p = 0.569). This suggests that Avoiders showed a faster reduction in freezing behavior during extinction than their Yoked counterparts. However, it is important to note that this effect was evident in the last of the three experiments only.

Similarly, suppression of lever pressing significantly decreased over extinction sessions in Experiment 1 (Q(3, 10.61) = 135.78, p < 0.001), Experiment 2 (Q(3, 10.98) = 258.58, p < 0.001) and Experiment 3 (Q(1, 9.45) = 14.77, p = 0.004). In Experiment 1, there was a significant interaction effect (Q(3, 10.61) = 7.14, p = 0.007, η_p_^2^ = 0.13) and an analysis of the simple main group effect on each day was performed, revealing that Avoiders showed more suppression than Yoked animals on day 1 (V = 135, p < 0.001) and less than Yoked animals on day 2 (V = 32.5, p = 0.024), suggesting steeper extinction learning in the Avoiders. Group differences leveled off by day 3. A fifth extinction session, with the platform now present, was included for both Avoider and Yoked groups in Experiment 1 (see results in Table S4 and Figure S4).

We hypothesized that, in the absence of the possibility to avoid, animals with a history of avoidance (group A) would show faster extinction learning than contingency-matched controls (group Y) in Experiment 1, but this was not clearly confirmed by our data. In Experiment 2, we hypothesized that the continued presence of the platform during extinction training would hinder extinction learning in group A, but this hypothesis was not supported by the data either. Finally, also in Experiment 3, we did not find clear differences between animals with and without a history of avoidance in speed of extinction learning.

### A history of avoidance partially counters reinstatement of fear after extinction

In Experiments 2 and 3, 24 h after the last extinction day (extinction day 4 in Experiment 2, extinction day 2 in Experiment 3), a reinstatement session took place which was followed by a reinstatement test (Fig 3, Table 3).

**Fig 3.**
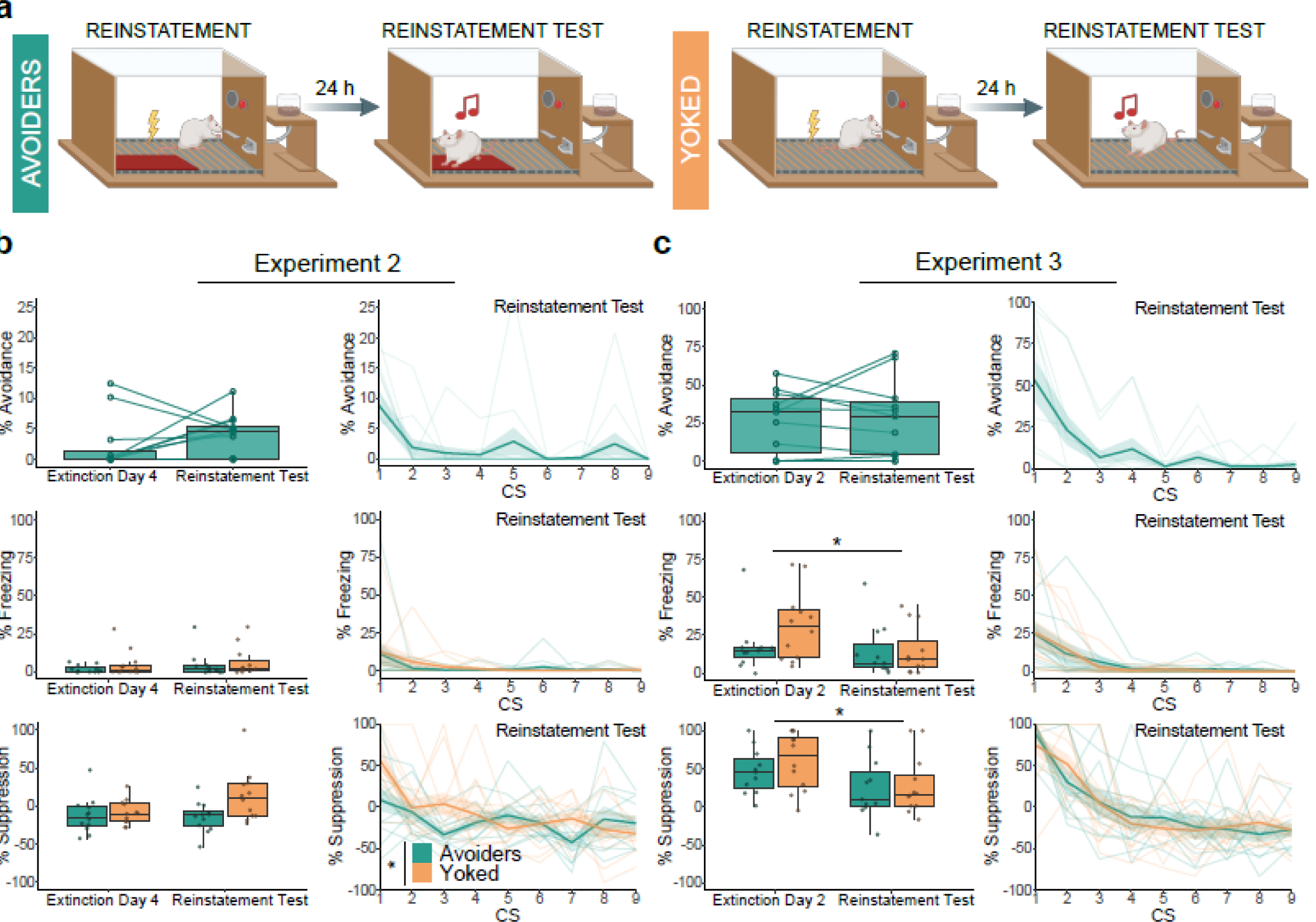
Reinstatement test results for Experiment 2 and 3. The box plots represent the average of the first 3 CSs on each day. The bold lines in the trial-by-trial plots represent the mean and the surrounding shaded area the standard error of the mean. Results are expressed in % of time during CS presentations. **a.** Graphical representation of reinstatement and the reinstatement test sessions in Experiment 2 and 3. **b.** Suppression of lever pressing differed significantly between the Avoider and the Yoked group (Q(1, 13.94) = 5.58, p = 0.042, η_p_^2^ = 0.175). **c.** Freezing decreased significantly between last day of extinction training and reinstatement test (Q(1, 11.33) = 6.72, p = 0.024, η_p_^2^ = 0.286), as did suppression of lever pressing (F(1, 21) = 5.31, p = 0.031, η_p_^2^ = 0.202). *p□<□0.05.

**Table 3.** Reinstatement test.

| Experiment | Measure | Statistical Test | Result | Effect size |
| --- | --- | --- | --- | --- |
| 2 | Avoidance | Wilcoxon signed ranks test | $V = 11, p = 0.363$ | $r = 0.348$ |
| | Freezing | Non-parametric mixed ANOVA | Group: $Q(1, 11.51) = 0.94, p = 0.353$<br>Day: $Q(1, 9.24) = 1.89, p = 0.202$<br>G*D: $Q(1, 9.24) = 0.27, p = 0.613$ | Group: $\eta_p^2 = 0.04$<br>Day: $\eta_p^2 = 0.142$<br>G*D: $\eta_p^2 = 0.005$ |
| | Suppression of lever pressing | Non-parametric mixed ANOVA | Group: $Q(1, 13.94) = 5.58, p = 0.042$<br>Day: $Q(1, 12.593) = 1.90, p = 0.191$<br>G*D: $Q(1, 12.593) = 1.93, p = 0.188$ | Group: $\eta_p^2 = 0.175$<br>Day: $\eta_p^2 = 0.087$<br>G*D: $\eta_p^2 = 0.131$ |
| | Rearing events | Non-parametric mixed ANOVA | Group: $Q(1, 8.89) = 0.36, p = 0.565$<br>Day: $Q(1, 10.84) = 1.68, p = 0.222$<br>G*D: $Q(1, 10.84) = 1.68, p = 0.222$ | Group: $\eta_p^2 = 0.055$<br>Day: $\eta_p^2 = 0.098$<br>G*D: $\eta_p^2 = 0.162$ |
| | Rearing duration | Non-parametric mixed ANOVA | Group: $Q(1, 8.23) = 0.48, p = 0.508$<br>Day: $Q(1, 8.33) = 2.02, p = 0.191$<br>G*D: $Q(1, 8.33) = 0.73, p = 0.418$ | Group: $\eta_p^2 = 0.081$<br>Day: $\eta_p^2 = 0.105$<br>G*D: $\eta_p^2 = 0.105$ |
| 3 | Avoidance | Paired t-test | $t(10) = -0.27, p = 0.792$ | $d = -0.081$ |
| | Freezing | Non-parametric mixed ANOVA | Group: $Q(1, 8.98) = 1.31, p = 0.283$<br>Day: $Q(1, 11.33) = 6.72, p = 0.024$<br>G*D: $Q(1, 11.33) = 2.25, p = 0.161$ | Group: $\eta_p^2 = 0.042$<br>Day: $\eta_p^2 = 0.286$<br>G*D: $\eta_p^2 = 0.161$ |
| | Suppression of lever pressing | Mixed ANOVA | Group: $F(1, 21) = 0.61, p = 0.443$<br>Day: $F(1, 21) = 5.31, p = 0.031$<br>G*D: $F(1, 21) = 0.23, p = 0.639$ | Group: $\eta_p^2 = 0.028$<br>Day: $\eta_p^2 = 0.202$<br>G*D: $\eta_p^2 = 0.011$ |
| | Rearing events | Non-parametric mixed ANOVA | Group: $Q(1, 9.06) = 1.23, p = 0.296$<br>Day: $Q(1, 11.67) = 3.05, p = 0.107$<br>G*D: $Q(1, 11.67) = 1.83, p = 0.201$ | Group: $\eta_p^2 = 0.051$<br>Day: $\eta_p^2 = 0.120$<br>G*D: $\eta_p^2 = 0.079$ |
| | Rearing duration | Non-parametric mixed ANOVA | Group: $Q(1, 11.06) = 1.04, p = 0.329$<br>Day: $Q(1, 11.97) = 1.44, p = 0.253$<br>G*D: $Q(1, 11.97) = 0.05, p = 0.823$ | Group: $\eta_p^2 = 0.098$<br>Day: $\eta_p^2 = 0.009$<br>G*D: $\eta_p^2 = 0.006$ |

Our various read-outs yielded partially different results. In Experiment 2, we observed a significant difference in the suppression of lever pressing between Avoiders and their Yoked counterparts (Q(1, 13.94) = 5.58, p = 0.042), with Yoked animals showing more suppression than Avoiders in the reinstatement test. However, for freezing or rearing we did not see any significant effects of group or day, nor did we see a significant increase in avoidance behavior in group A (see Table 3). In sum, to the extent that reinstatement was obtained at all in Experiment 2, it was restricted to Yoked controls but absent in Avoiders.

Surprisingly, in Experiment 3, the reinstatement session did not reinstate fear in either of the groups, even though the foot shock administered for reinstatement had double the intensity of that of Experiment 2 (0.8 mA versus 0.4 mA). Against our expectations, we even saw a significant decrease in freezing (Q(1, 11.33) = 6.72, p = 0.024) and in suppression of lever pressing (F(1, 21) = 5.31, p = 0.031) from the last extinction session to the reinstatement test. No significant differences were found in avoidance behavior, nor in rearing events or duration.

There are a number of important differences between these two experiments. Animals in Experiment 2 went through 4 sessions of extinction while animals in Experiment 3 had only 2 extinction sessions. Also, animals in Experiment 3 had a classical conditioning session (without platform) at the very start of the experiment and received a reinstatement shock of a higher intensity. However, none of those differences readily explain why reinstatement was even less prominent in Experiment 3 than in Experiment 2. An inspection of the reinstatement session’s video footage of both experiments showed that rats presented clear unconditioned responses to the shocks, ruling out an explanation in terms of equipment failure.

A priori, for Experiment 2, we expected that Avoider animals would show more return of fear than Yoked animals in the reinstatement test session, owing to the fact that the presence of the platform would reduce extinction learning, rendering fear more easily reinstated. Not only did we not see differences between groups in the majority of behavioral read-outs, we even found that Yoked controls suppressed lever pressing more than Avoider animals in the reinstatement test. In Experiment 3, no significant differences between groups were observed altogether.

### Spontaneous recovery is unaffected by a history of avoidance

In Experiments 2 and 3, after reinstatement testing, we performed a platform switch session (i.e., rats in group A were now tested without the platform, whereas rats in group Y were now tested with the platform present), which in Experiment 2 was followed by an additional session using a non-grid floor in both groups. No group differences were found in any of these sessions, as discussed in the Supplement (Table S6). In Experiment 3, a further test session was followed by a delayed test session one week later to assess spontaneous recovery (Fig 4, Table 4).

**Fig 4.**
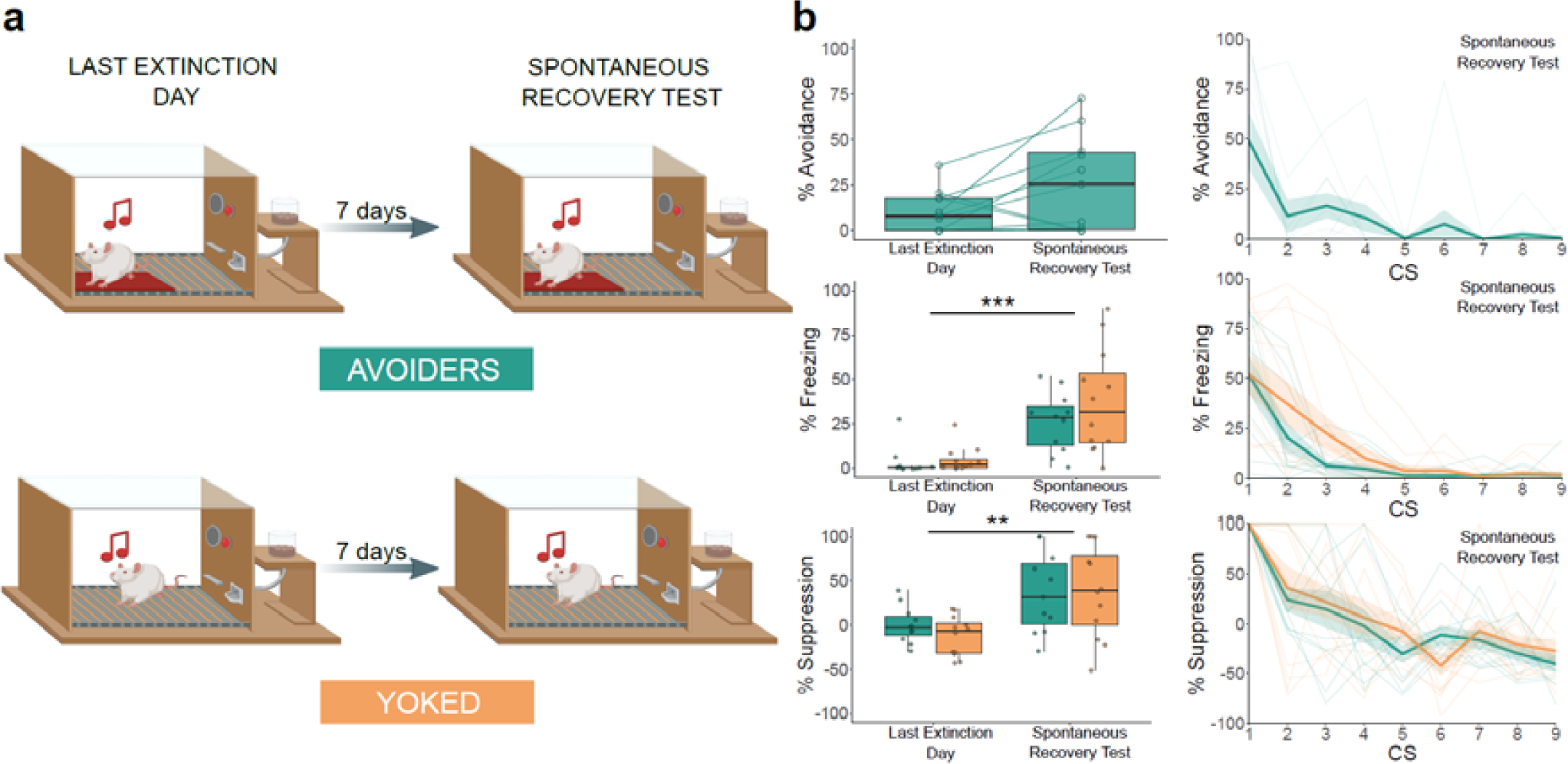
Spontaneous recovery test results for Experiment 3. The box plots represent the average of the first 3 CSs on each day. The bold lines in the trial-by-trial plots represent the mean and the surrounding shaded area the standard error of the mean. Results are expressed in % of time during CS presentations. **a.** Graphical representation of the last extinction session and the spontaneous recovery test session. **b.** Freezing increased significantly from the last extinction session to the spontaneous recovery test session (Q(1, 9.72) = 25.61, p < 0.001, η_p_^2^ = 0.600). Similarly, suppression of lever pressing increased significantly between the last extinction session and the spontaneous recovery test session (Q(1, 10.65) = 10.99, p = 0.007, η_p_^2^ = 0.463). ** p < 0.01, ***p□<□0.001.

**Table 4.**
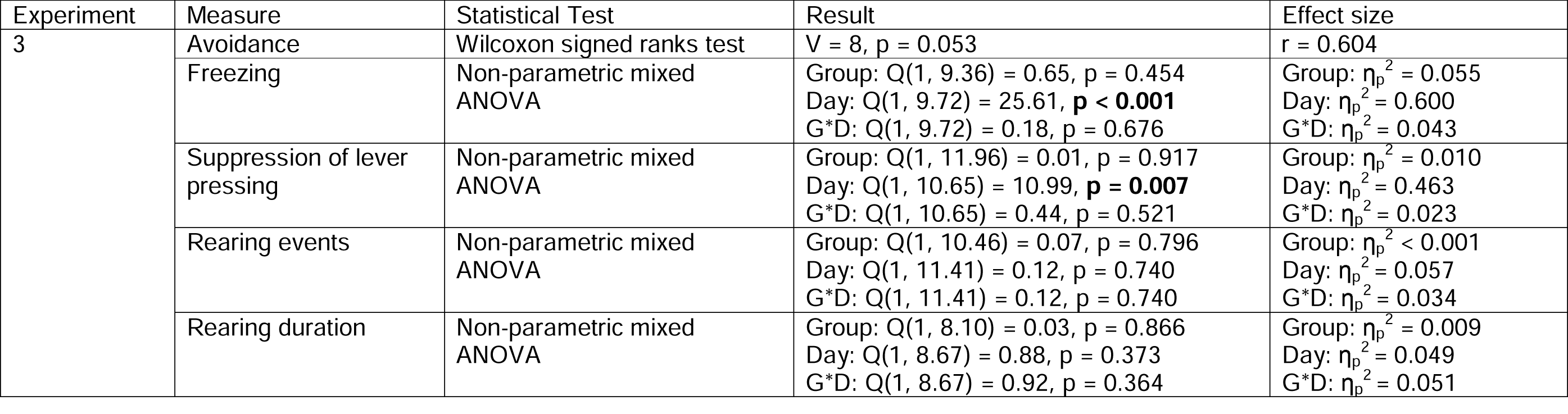
Spontaneous Recovery test.

| Experiment | Measure | Statistical Test | Result | Effect size |
| --- | --- | --- | --- | --- |
| 3 | Avoidance | Wilcoxon signed ranks test | $V = 8, p = 0.053$ | $r = 0.604$ |
| | Freezing | Non-parametric mixed ANOVA | Group: $Q(1, 9.36) = 0.65, p = 0.454$<br>Day: $Q(1, 9.72) = 25.61, p < \mathbf{0.001}$<br>G*D: $Q(1, 9.72) = 0.18, p = 0.676$ | Group: $\eta_p^2 = 0.055$<br>Day: $\eta_p^2 = 0.600$<br>G*D: $\eta_p^2 = 0.043$ |
| | Suppression of lever pressing | Non-parametric mixed ANOVA | Group: $Q(1, 11.96) = 0.01, p = 0.917$<br>Day: $Q(1, 10.65) = 10.99, p = \mathbf{0.007}$<br>G*D: $Q(1, 10.65) = 0.44, p = 0.521$ | Group: $\eta_p^2 = 0.010$<br>Day: $\eta_p^2 = 0.463$<br>G*D: $\eta_p^2 = 0.023$ |
| | Rearing events | Non-parametric mixed ANOVA | Group: $Q(1, 10.46) = 0.07, p = 0.796$<br>Day: $Q(1, 11.41) = 0.12, p = 0.740$<br>G*D: $Q(1, 11.41) = 0.12, p = 0.740$ | Group: $\eta_p^2 < 0.001$<br>Day: $\eta_p^2 = 0.057$<br>G*D: $\eta_p^2 = 0.034$ |
| | Rearing duration | Non-parametric mixed ANOVA | Group: $Q(1, 8.10) = 0.03, p = 0.866$<br>Day: $Q(1, 8.67) = 0.88, p = 0.373$<br>G*D: $Q(1, 8.67) = 0.92, p = 0.364$ | Group: $\eta_p^2 = 0.009$<br>Day: $\eta_p^2 = 0.049$<br>G*D: $\eta_p^2 = 0.051$ |

The comparison of the delayed test session in Experiment 3 to the preceding one indicates that there was spontaneous recovery in both groups, as evidenced by increases in freezing (Q(1, 9.72) = 25.61, p < 0.001) and in suppression of lever pressing (Q(1, 10.65) = 10.99, p = 0.007). There were no significant group differences. Avoidance behavior in group A was higher during the spontaneous recovery test than during the preceding session, but this increase did not reach significance.

In sum, in contrast to our initial hypothesis, Yoked animals did not show a different sensitivity to spontaneous recovery of fear as indexed through freezing and suppression of lever pressing than Avoiders.

### Persistence of avoidance

Persistence of avoidance (i.e., high residual rates of avoidance) despite extinction^14^ or extinction with response-prevention^15^ has previously been reported in approximately 25% of rats in the platform-mediated avoidance task. We evaluated the persistence of avoidance in Experiments 2 and 3, where the platform was present during extinction training in group A.

In Experiment 2, we classified a rat as a persistent Avoider if on the third extinction day, during the first block, its time on the platform during the CS was more than 50%. We did not identify any rats meeting that criterion, so all rats in Experiment 2 were considered non-persistent Avoiders according to our preregistered definition. We reanalyzed the data applying criteria reported in previous reports^14,15^, but still failed to detect any persistent Avoiders.

In Experiment 3, given the reduction from four to two days of extinction, we assessed resistant rather than persistent avoidance. As preregistered, rats were classified as resistant Avoiders if they spent more than 50% of the duration of CS presentation on the platform across the first block of the reinstatement test or the first block of the spontaneous recovery test. We identified two rats that qualified as resistant Avoiders (2/11: 18%). When applying the criteria used in a previous study^15^, we found only 18% of resistant Avoiders as well. Only one rat out of the two was identified consistently as resistant Avoider using both criteria. With regards to performance during the reinstatement test, rats could be divided in three clusters, from 0 to 25% of avoidance (5 rats), between 25% and 50% (4 rats), and between 50% and 75% (2 rats). For the spontaneous recovery test, we similarly find 3 clusters, ranging from 0 to 10% of avoidance (5 rats), 50 to 75% of avoidance (4 rats), and more than 75% of avoidance (2 rats).

The preregistered comparisons between persistent and non-persistent Avoiders or resistant versus non-resistant Avoiders were not performed due to the low number of persistent and resistant Avoiders.

## Discussion

Here we showed that a history of avoidance has no clear effect on the subsequent extinction of auditory cued fear in a platform-mediated avoidance procedure that constitutes a realistic approach/avoidance conflict in rats. Regardless of whether the possibility for avoidance was maintained during extinction (Experiment 2 and 3) or not (Experiment 1), no marked differences in extinction learning were found between rats that were previously allowed to engage in avoidance and their Yoked controls. We also investigated the return of fear in rats with a history of avoidance compared to their Yoked counterparts. In reinstatement, we observed less return of lever press suppression in rats allowed to avoid than in the Yoked rats in Experiment 2 but not in Experiment 3. No other significant differences in reinstatement were observed. Experiment 3 further yielded a comparable degree of spontaneous recovery in both groups. An overview of the main results can be found in Table 5.

**Table 5.**
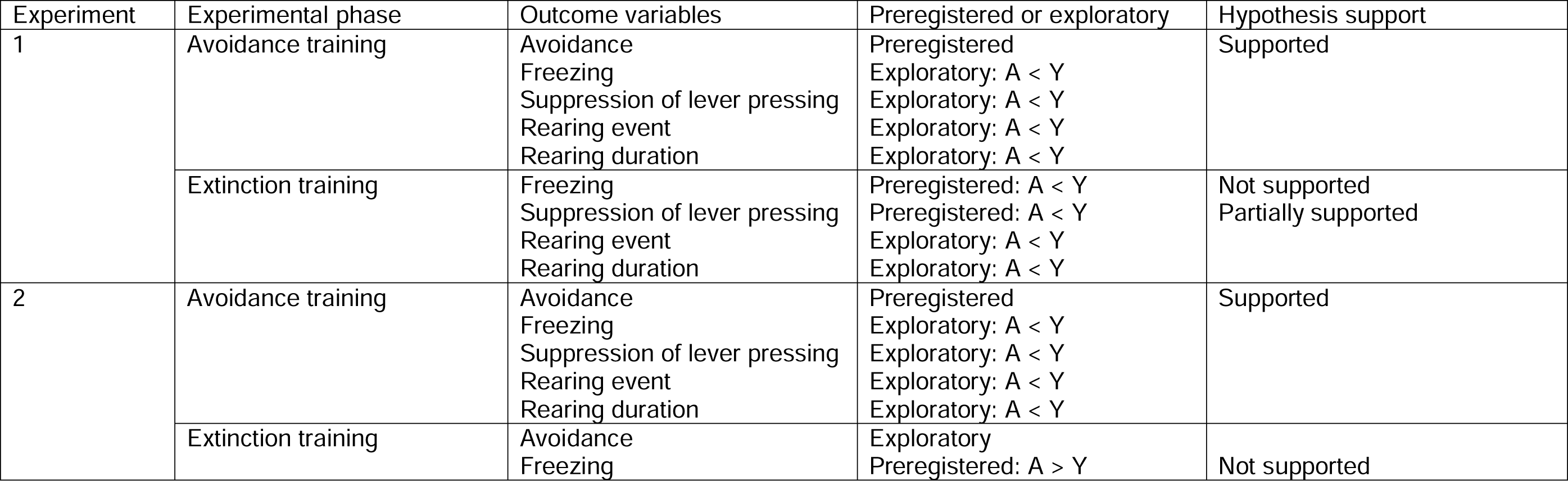
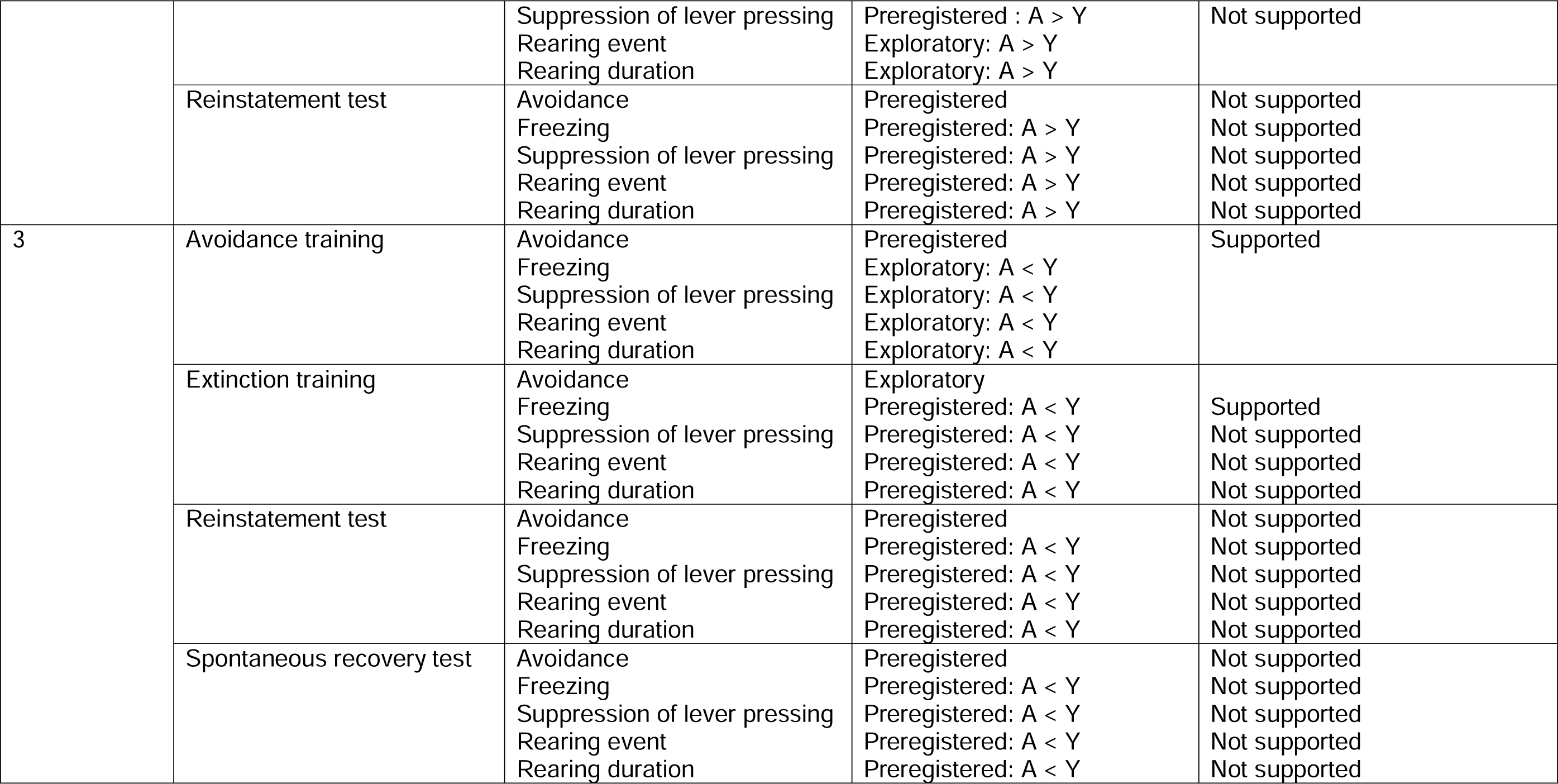
Results summary main hypotheses.

Across the three experiments reported here, we assessed four different behavioral read-outs (avoidance, freezing, suppression of lever pressing and rearing). As mentioned in the Results section, the various measures did not always yield similar outcomes. For instance, in Experiment 2, when assessing reinstatement (Fig 3), we observed a significant increase in suppression of lever pressing, no increase in freezing, and a nominal increase in avoidance that was non-significant. So, while our measured outcomes are related and mostly convergent, they are not identical.

Studies have shown a high correlation between freezing and suppression of lever pressing^20^, which makes sense given that freezing by definition implies the cessation of movement and, therefore, lever pressing. Additionally, lesions to the lateral amygdala^21^ and the central nucleus of the amygdala^22,23^ block freezing as well as suppression of lever pressing and suppression of licking. However, freezing is by no means necessary to generate suppression of lever pressing. Lesions to the periacqueductal gray after fear conditioning block freezing but not suppression of lever pressing^24^ and chemogenetic inhibition of cells in the ventrolateral periacqueductal gray has no effects on fear acquisition but impairs extinction of suppression of lever pressing independently of freezing^25^. Therefore, even if these two behaviors are fear-related, they are different and worth investigating.

Regarding comparisons of freezing and avoidance, in the PMA task, these behaviors are not incompatible because rats can freeze on the platform. This is different from other active avoidance procedures, such as the two-way active avoidance task, where freezing is inversely correlated with avoidance responding, with poor avoiders showing high degrees of freezing, and lesions in the central amygdala reversing poor avoidance^26^. In the present PMA procedure, in contrast, we observed a significant increase of avoidance as well as freezing during avoidance training in Experiments 1 and 2. This was not the case in Experiment 3, where avoidance training was preceded by a session of purely Pavlovian conditioning and where we observed a significant increase in avoidance only. Our results may seem to contradict results from two-way avoidance tasks where a reduction of freezing is typically observed as avoidance training progresses and avoidance increases^27^, something that has also been observed in previous reports using PMA^13^. However, in our study, avoidance training lasted for just two days, which is sufficient to obtain asymptotic avoidance but likely insufficient to obtain a reduction in freezing. Additionally, other authors that used the PMA task have reported persistent avoidance in around 25% of the animals^14,15^, while in our case the number of persistent avoiders was considerably lower. One of the reasons for this difference may again be the more limited duration of avoidance training in our studies, which lasted only two days, as opposed to the ten-day training period reported by other researchers. Despite our rats reaching similar rates of avoidance as in previous research, it may be that our training was not sufficiently long to result in persistent responding during extinction. Due to our Yoked design, increasing the duration of avoidance training was not viable because Yoked rats would have prematurely entered extinction training due to the lack of foot shocks earned by their corresponding animals in group A when avoidance reaches asymptote.

The results reported here are surprising given the considerable amount of data regarding the effects of ongoing avoidance on the efficacy of extinction training^4,5,28^. However, in most prior work, avoidance responses have little to no cost (i.e., shuttling requires little caloric expenditure). In experiments with human participants, researchers have found that when avoidance has a clear cost, participants show less avoidance behavior and learn extinction more readily^29^. In human research, one such cost can be an increased time delay, meaning longer trials and extended experiment duration, which is reported to be aversive^30,31^, whereas other studies have used monetary losses as response cost^32,33^. In our case, if rats step onto the platform to avoid foot shock, they miss out on the opportunity to obtain food rewards that are highly desirable because the animals have limited access to food during the experiment. This realistic cost of avoidance responding does not impact its acquisition in the face of threat, but we speculate that it may prevent the adverse effects that avoidance has been thought to have on extinction in previous studies using lever press avoidance^34^, shuttle avoidance^5^ and by similar tasks^35^. In other words, if avoidance has a sufficient cost associated with it, it will reduce readily under extinction and have no negative consequences for the extinction of other fear responses. This would mean that, in a clinical setting, avoidance behaviors associated with a sufficient cost are not worrisome. At the same time, we have not found clear evidence for a beneficial effect of a history of avoidance on extinction learning or retention either, apart from a (very) partial reduction in return of fear. Thus, neither do we have reasons to encourage avoidance behaviors.

One limitation of our study is that the obtained results apply to male rats only. It has been reported that female rats acquire active avoidance faster than males^36^, even when controlling for weight and age differences^37^. A study on the extinction of avoidance, using a passive avoidance task, found that both sexes showed similar freezing during the last minute of acquisition, similar avoidance but females showed reduced freezing during extinction^38^. Finally, a recent study employing the PMA task reported that female rats were more resistant to the extinction of avoidance than their male counterparts^39^. Therefore, it would be worthwhile to investigate if an experiment using our design yields different results when using females instead of male rats.

Another limitation of the experiments presented here could be the limited length of avoidance training (although at the end of the two days, last 3 trials, rats did spend 70-75% of the CS duration on the platform/reached an asymptote). Additional avoidance training sessions could reduce freezing to lower levels and possibly increase the chances of finding persistent Avoider rats. However, our Yoked design makes it difficult to increase the avoidance training sessions, and modifications in that regard should be made to avoid that the Yoked group receives what would functionally be extinction training, rendering comparisons between Avoiders and their Yoked controls exceedingly complex.

Finally, it could be interesting to replicate the current studies using Wistar-Kyoto rats. In the lever-press avoidance task, this strain showed facilitated avoidance acquisition^40,41^ and deficits in avoidance extinction^42^ compared to Sprague Dawley rats. Wistar-Kyoto rats are a rat model of anxiety-like behavior and, given that anxiety patients engage in avoidance responses despite their high cost and sometimes persist in costly avoidance in the absence of threat^32^, it would be worthwhile to test if the Wistar-Kyoto strain could model this phenotype in the PMA task.

In summary, this study examined the effects of avoidance history on subsequent extinction of auditory cued fear in a platform-mediated avoidance procedure. We reported that two days were sufficient for rats to acquire avoidance and we showed that extinction of fear occurred similarly in rats with or without a history of avoidance. Additionally, when the platform remained present during extinction, rats still extinguished avoidance and fear similarly regardless of their history of avoidance. We additionally observed partial effects on reinstatement of fear, where Yoked animals showed higher return of lever press suppression than Avoiders, but we did not observe any differences in spontaneous recovery.

## Methods

### Preregistration and data availability

All experiments, sample sizes, and analysis plans were registered on the Open Science Framework (OSF) before the start of data collection (Experiment 1: https://osf.io/acngu/; Experiment 2: https://osf.io/y5p3r/; Experiment 3: https://osf.io/2kbg5/). Data and scripts for every experiment can also be found there.

### Subjects

All experiments were performed in accordance with Belgian and European laws (Belgian Royal Decree of 29/05/2013 and European Directive 2010/63/EU) and the ARRIVE 2.0 guidelines^43^ and approved by the KU Leuven animal ethics committee (project license number: 011/2019). The number of animals used was calculated according to previous studies. All the studies were performed in 8-week-old male Sprague Dawley rats (270-300 g at arrival; Janvier Labs, Le Genest_-_Saint_-_Isle, France). Animals were housed in groups of 3 on a 12-hour light-dark cycle (lights on at 7am) and experiments were performed between 10am and 4pm. Cages had bedding and cage enrichment in the form of a red polycarbonate tunnel hanging from the top of the grid. Water was available ad libitum for the whole experiment, except during behavioral testing. Food was provided ad libitum until one day before the start of the experiments. From then on, rats were fed food ad libitum for an hour after each test session. Animals were habituated to handling for two days before the start of the experiment.

### Apparatus

Six identical operant chambers (30.5 cm width, 25.4 cm depth and 30.5 cm height; Rat Test Cage, Coulbourn Instruments, Pennsylvania, USA) were used simultaneously and were enclosed in sound-attenuating boxes. A 12.8 cm by 15.2 cm platform made of a non-transparent red plastic sheet was used in some parts of the experiments. It covered about ¼ of the grid floor, and it was always placed in the corner opposite to the lever. Experiments were run with the house light off and behavior was recorded with an infrared IP camera (Foscam C1, Shenzhen, China).

### Lever press training

Lever press training consisted of an hour of training for ten consecutive days. Rats were trained to lever press for a grain-based pellet (45 mg 5TUM, TestDiet, St. Louis, MO, USA) on a variable interval schedule of reinforcement averaging 30 seconds (VI-30 s). Rats that did not reach the criterion of an average of 15 lever presses per minute across the VI-30 s session were excluded from further analyses (rat A0 from Experiment 3).

The first training phase consisted of a full session of magazine training with the lever retracted, where food pellets were delivered at fixed 2-min intervals. The second phase of lever press training consisted of magazine training with the lever extended, where food pellets were delivered at fixed 2-min intervals, supplemented with direct delivery of a food pellet upon each lever press. This second phase lasted one day minimally, and required at least 1 lever press/min on average to pass to the next phase. If there were no lever presses during the first half of the third day of this second phase of training, hand-shaping was performed. The next phases consisted of lever press training on increasing variable ratio (VR) schedules, progressing from VR 3 through VR 5 and VR 15 to VR 30, where pellets were delivered after a variable number of lever presses, with an increasing average reinforcement criterion per session. In VR 3, a criterion of at least 3 presses per minute was set to pass to the next schedule, a criterion of at least 5 lever presses per minute for VR 5, a criterion of at least 10 lever presses per minute for VR 15, and a criterion of at least 15 lever presses per minute for VR 30. The last phase consisted of training on a VI 30 schedule where the first lever press after a variable refractory period averaging 30 seconds yielded pellet delivery.

### Experiment 1

Rats (n=24) were divided into two equal groups: group A (Avoiders) and group Y (Yoked) (Fig 5a). Avoidance training lasted for two days. On each day, nine CSs (a pure tone of 3 kHz) of 30 s were presented, with an ITI averaging 180 s (from 150 s to 210 s). Each CS co-terminated with a foot shock US of 2 s and an intensity of 0.4 mA. Avoider rats had the possibility to avoid the US by stepping onto a platform, whereas Yoked rats did not have a platform present in the box. The number of shocks and the length of shocks received by the Avoider animals (A) was scored to allow the presentation of the same CS-US contingency to animals in the Yoked group (Y). Yoked rats always received the first scheduled CS-US pairing of the session. However, for the remaining trials, the CS-US contingency depended on what their companion animal in the A group did: (1) if A had avoided the shock altogether, Y received the CS without a US, (2) if A had escaped to the platform during the US and thus managed to avoid part of it, Y received the CS with a US of 1 s duration on the first day of conditioning or 0.5 s on the second day of conditioning, (3) if A had not stepped onto the platform at any time during the US, Y received a regular CS-US pairing with a US of 2 s. To achieve this, we used a staggered onset, meaning that Avoider rats were trained one day ahead of their Yoked counterparts. We manually scored if and for how long each of the 9 USs were received by the Avoider animal and then programmed the new tasks for each Yoked counterpart accordingly.

**Fig 5.**
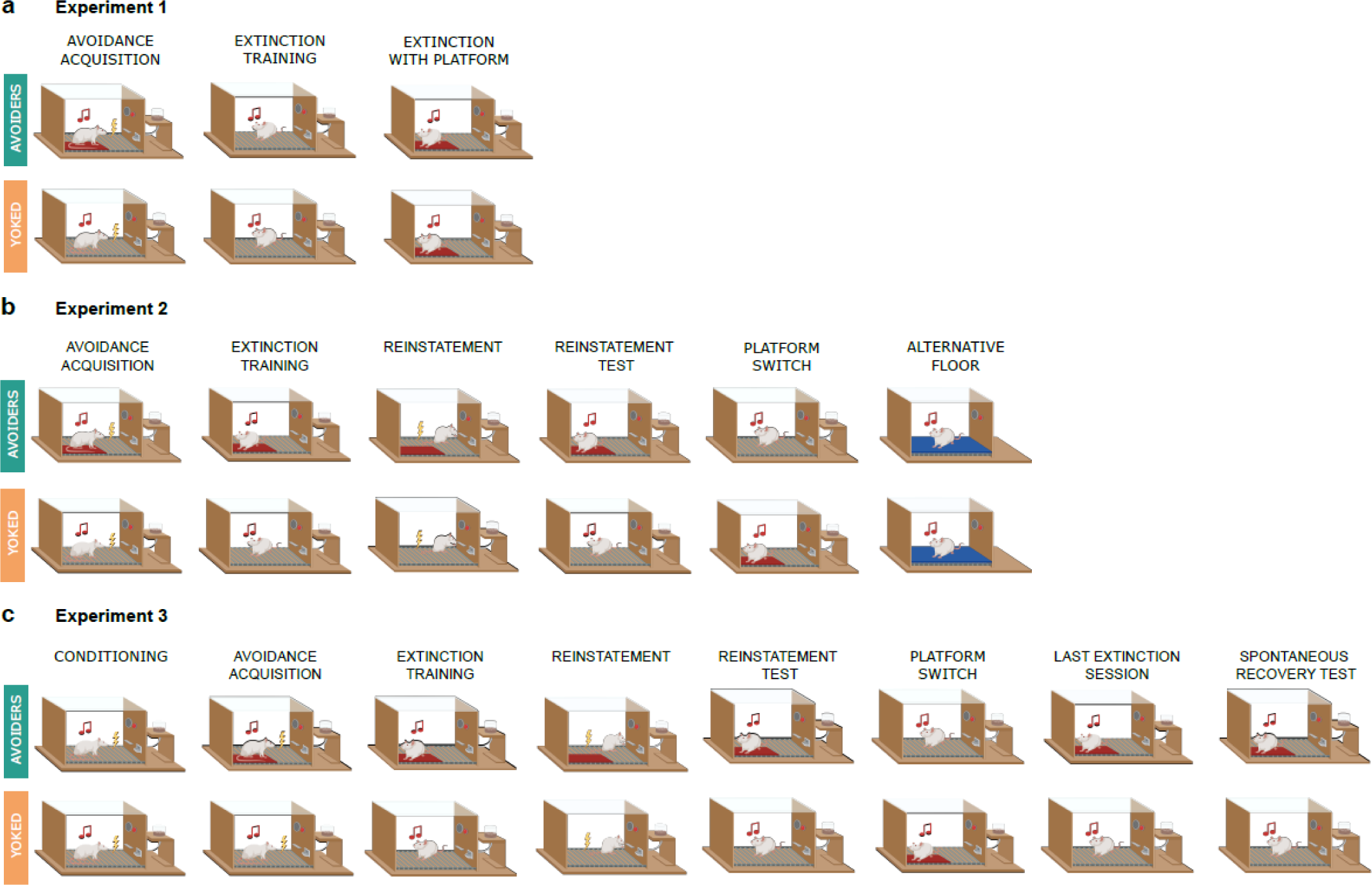
Graphic representation of Experiments 1-3. **a.** Graphic representation of the different phases of Experiment 1. **b.** Graphic representation of the different phases of Experiment 2. **c.** Graphic representation of the different phases of Experiment 3.

After the two days of avoidance training, rats went through extinction training for four days, with the platform absent in both groups, where nine CSs of the same characteristics as above were presented per day, without foot shock. Finally, a fifth extinction day with nine CSs was conducted with the platform present for both groups.

### Experiment 2

Rats (n=24) were divided into two equal groups: group Avoiders (A) and group Yoked (Y) (Fig 5b). Experiment 2 had the same avoidance and extinction training as Experiment 1 with the exception that group A had a platform present during the four extinction sessions. By adding the platform during extinction for this group, we wanted to investigate whether animals in group A would keep exhibiting avoidance despite the absence of USs during extinction. This thus allowed us to specifically test if avoidance during extinction training defied extinction learning as previously reported^4^.

Additionally, Experiment 2 had a reinstatement session where, after 3 minutes of habituation, an unsignaled foot shock of 1 s and 0.4 mA was delivered. The following day, a reinstatement test was performed where the setup was the same as during extinction training, including the presence of the platform for group A but not group Y. The reinstatement session and test were added to investigate any differences in return of fear between group A and group Y. The next day, a platform switch session took place. The setup was the same as for the preceding extinction sessions with the difference that the platform was removed for group A and introduced for group Y. This session was added to the experiment to assess if the lack of platform, and therefore change of context, could increase fear in group A. An additional extinction session followed the platform switch session, with the platform again present for group A and absent for group Y. The last session consisted of an alternative floor test, where a floor was made from a plastic placemat (Hema, Belgium) coated with fake leather fabric (de Banier, Belgium) that fitted on top of the entire grid floor of the box, and the lever was retracted leaving freezing as the only outcome measure. This session was added to evaluate freezing to the CS in a new context and without any interference of lever-pressing.

Due to technical issues during data collection, the videos from 4 rats of group Y during the second extinction day and videos from 4 rats of group A during the third extinction day were lost. Given the nature of the missing values, it is safe to assume that the missing data would be completely at random, therefore we substituted the missing data by group averages for avoidance, freezing and rearing events and percentage in those cases^44^.

### Experiment 3

First, all rats (n=24) went through a single classical conditioning session, without platform, during which nine CSs co-terminated with a foot shock of 1s and 0.4 mA (Fig 5c). Then, as in previous experiments, rats were divided into two equal groups: group Avoiders (A) and group Yoked (Y). Experiment 3 had the same avoidance and extinction sessions as Experiment 2, with the exception that there were only two extinction sessions instead of four. Extinction was shortened in this experiment to assess if a shorter extinction training would affect later return of fear. Experiment 3 also included a similar reinstatement session as in Experiment 2, be it that the reinstatement shock had an intensity of 0.8 mA instead of 0.4 mA. This change in intensity was implemented to produce stronger reinstatement of fear. The experiment followed the design of Experiment 2, with a reinstatement test and platform switch session. On the next day, one last extinction session was performed, with the same characteristics as described previously, and with the main aim of providing a baseline for comparison with the spontaneous recovery test presented one week later. This experiment thus contained two return-of-fear tests (reinstatement and spontaneous recovery) to thoroughly evaluate return of fear and avoidance in both groups. One rat from group A was excluded because it did not reach lever press criteria.

### Data analysis

Behavior during CS presentations was scored manually from videos. The experimenter was blinded for test session and group whenever possible (platform presence gave away group allocation in some test sessions). Freezing, avoidance and rearing were scored as percentage of CS duration. Avoidance was considered to be present whenever at least two paws of the animal were on the platform and the center of mass of the rat was over the platform as well. Freezing was defined as full immobility except for the minimal movements associated with breathing. A rat was considered to be rearing when it lifted two paws from the floor; no distinction between supported or unsupported rearing was made. The rearing event stopped once one of the front paws touched the floor or the lever. We scored the number of rearing events during a CS as well as the percentage of time spent rearing during a CS presentation.

Lever presses produced one minute before the CS (pre-CS) and during the CS were aggregated per block (each block consists of 3 CSs), after which the following formula was applied to calculate suppression of lever pressing: (pretone rate - tone rate)/(pretone rate + tone rate) x 100 (Quirk et al., 2000). The values ranged from no suppression of lever pressing (0%) to full suppression of lever pressing (100%). For blocks where pretone and tone values were both 0, the rat’s suppression was substituted by the mean group average for that block.

Most of the statistical analyses were preregistered on OSF (for details and deviations, see below). The 9 CSs were grouped in blocks of 3 CSs. The blocks of the experimental groups (Avoiders vs Yoked) were compared using mixed repeated measures (RM) ANOVA and, if significance was reached, followed by post-hoc tests with multiple-test corrections. Avoidance behavior within the Avoider group was evaluated using paired t-tests or one-way repeated-measures ANOVAs. When assumptions were violated, non-parametric statistical analyses were performed. For the RM mixed ANOVAs, we applied robust ANOVAs with a trimmed means approach^45^ using the WRS2 package^46^ from R (Version 4.1.3; R Core Team, 2022)^47^ and R Studio (Version 9.1.372; RStudio Team, 2021)^48^. Where a non-parametric mixed ANOVA was performed, we report the effect sizes as obtained from a parametric mixed ANOVA given that there are no clear guidelines for how to derive effect sizes for non-parametric mixed ANOVAs. All other statistical analyses were performed using afex^49^ or rstatix^50^. Data were processed and plotted using the tidyverse^51^, reshape^52^ and zoo^53^ packages.

#### Experiment 1

We preregistered between-group comparisons of freezing and of suppression of lever pressing during extinction days without platform with RM ANOVAs (Group x Day x Block). In addition, we preregistered separate analyses between groups to assess avoidance, freezing and suppression of lever pressing in the extinction test with the platform (RM ANOVA, Group x Block). In case of significant effects or interactions, post-hoc tests were performed. Note that we had preregistered an outlier analysis (ROUT test, GraphPad Prism) for suppression of lever pressing during the extinction sessions, but in the end decided to not exclude any potential outliers. As a secondary hypothesis we preregistered the comparison of avoidance behavior between day 1 and 2 using a one-sided dependent-samples t-test in the avoidance group only.

Analyses of rearing behavior were not part of the preregistered analysis plan for Experiment 1. Additionally, analyses comparing freezing and suppression of lever pressing between groups during avoidance learning were not preregistered and are therefore exploratory.

#### Experiment 2

We preregistered the same analyses as above for the extinction session, adding the analysis of rearing for this experiment. We also preregistered a between-group comparison of freezing, suppression of lever pressing and rearing during extinction session 4 and the reinstatement of fear test (RM ANOVA, Group x Day). In case of significant effects or interactions, post-hoc tests were performed. In the Avoider group, one-sided dependent-samples t-tests were used to compare time spent on the platform during CS presentation during extinction session 4 and the reinstatement of fear test. We also preregistered separate independent-samples t-test analyses to assess freezing, suppression of lever pressing and rearing in the platform switch session. A one-sided independent-samples t-test was preregistered to assess differences in freezing and rearing in the alternative floor session. We preregistered the comparison of avoidance behavior between day 1 and day 2, using a one-sided dependent-samples t-test.

Finally, we also preregistered a comparison between non-persistent and persistent Avoider rats. A rat would be considered a persistent Avoider when the time on the platform during the CS was more than 50% during the first block on the 3rd extinction day, a similar criterion as used in a previous study^14^. However, we did not have any persistent Avoider rats in this experiment and therefore could not perform this analysis.

#### Experiment 3

Here we preregistered the same analyses as reported for Experiment 2. Additionally, we preregistered a between-group comparison of freezing, suppression of bar pressing and rearing during the first block of the last extinction session and the spontaneous recovery test (RM ANOVA, Group x Day).

In addition, we conducted a Wilcoxon test that was not preregistered, to compare the percentage of avoidance between the first block of the last extinction day and the spontaneous recovery test.

## Data availability

The behavioral data and analysis scripts that support the findings of this study are available on OSF (Experiment 1: https://osf.io/acngu/; Experiment 2: https://osf.io/y5p3r/; Experiment 3: https://osf.io/2kbg5/). Raw videos are available upon reasonable request.

## Acknowledgements

This work was supported by KU Leuven Research Grant 3H190245 and FWO PhD fellowship 11K3821N. The funder played no role in study design, data collection, analysis and interpretation of data, or the writing of this manuscript. Graphical representations in the figures were created with BioRender.com.

## Author contributions

AL-M, LL and TB conceived and designed the study. AL-M collected and performed the data analysis. AL-M wrote the first draft of the manuscript. LL and TB edited and critically reviewed the manuscript. All authors reviewed the results and approved the final version of the manuscript.

