## Supplementary figures and tables for "A history of avoidance does not impact extinction learning in male rats"

Fig S1. Pavlovian phase Experiment 3

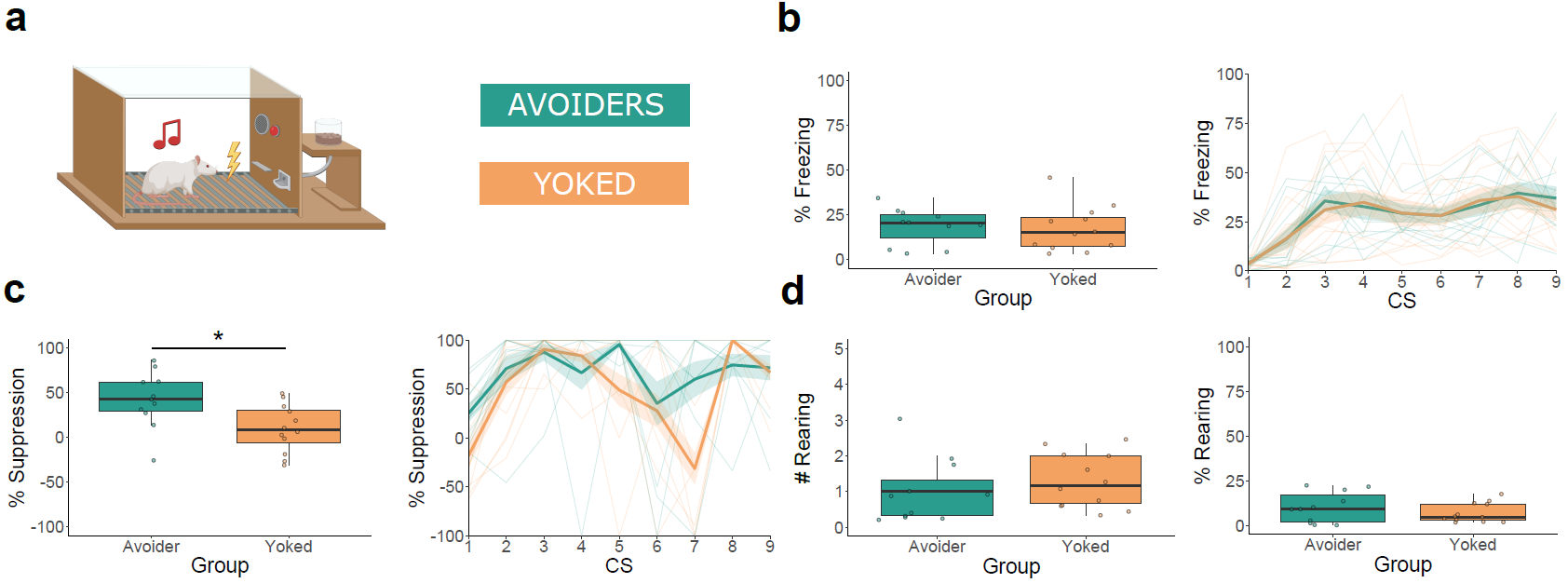

*Fig S1*. Pavlovian acquisition results for Experiment 3. The box plots represent the average of the first 3 CSs. The bold lines in the trial-by-trial plots represent the mean and the surrounding shaded area the standard error of the mean. Results are expressed in % of time during CS presentations, except in rearing behavior where the first plot expresses the average number of rearing bouts during CS presentations. **a.** Graphical representation of the purely Pavlovian session which was identical for both groups. **b.** Both groups show similar freezing in Pavlovian acquisition. **c.** Rats assigned to be Avoiders in the subsequent avoidance learning phase suppress lever pressing more than their Yoked counterparts (t(19.8) = 2.64, p = 0.016, d = 1.1). **d.** The rate and duration of rearing is similar across groups in Pavlovian acquisition. *p < 0.05.

Fig S2. Additional behaviors in avoidance training Experiment 1-3

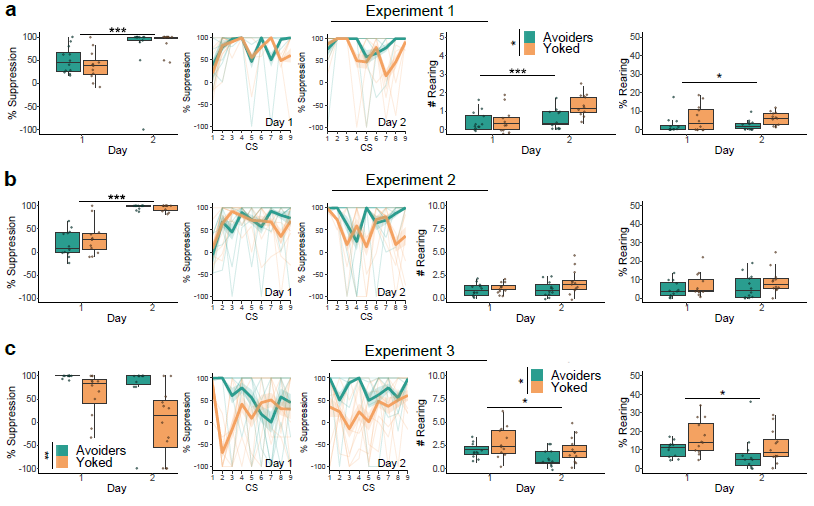

*Fig S2.* Additional behaviors in avoidance training for Experiments 1 -3. The box plots represent the average of the first 3 CSs of every day. The bold lines in the trial-by-trial plots represent the mean and the surrounding shaded area the standard error of the mean. Results are expressed in % of time during CS presentations, except in rearing behavior where the first plot expresses the average number of rearing bouts during CS presentations. **a.** Both groups increased their suppression of lever pressing (F(1, 22) = 18.32, p < 0.001, η_p_^2^ = 0.454) and rearing events (Q(1, 13.05) = 18.32, p < 0.001, η_p_^2^ = 0.281) across training days in Experiment 1. However, Yoked rats showed increased rates (Q(1, 13.43) = 4.95, p = 0.044, η_p_^2^ = 0.267) and duration of rearing compared to Avoiders (Q(1, 9.56) = 6.96, p = 0.026, η_p_^2^ = 0.234). **b.** Both groups increased their suppression of lever pressing (Q(1, 12.58) = 161.70, p < 0.001, η_p_^2^ = 0.854) across training days in Experiment 2. **c.** Avoider rats showed increased suppression of lever pressing across avoidance training compared to their Yoked counterparts (Q(1, 6.33) = 14.59, p = 0.008, η_p_^2^ = 0.38), but Yoked rats reared more often than Avoiders across avoidance training sessions (F(1, 21) = 4.68, p = 0.042, η_p_^2^ = 0.182) in Experiment 3. A decrease in amount (F(1, 21) = 6.66, p = 0.017, η_p_^2^ = 0.241) and duration of rearing (Q(1, 10.03) = 6.03, p = 0.034, η_p_^2^ = 0.105) across avoidance training session is observed in Experiment 3. *p < 0.05, ** p < 0.01, ***p < 0.001.

Fig S3. Rearing in extinction training Experiments 1-3

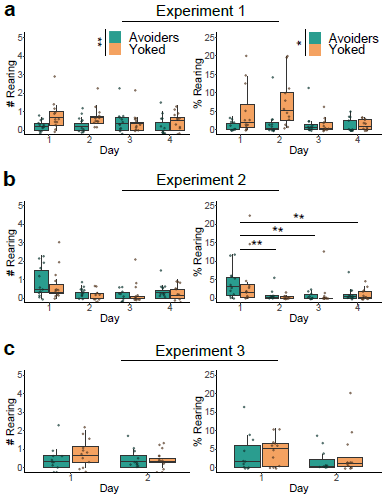

*Fig S3.* Rearing events and duration in extinction training Experiments 1-3. The box plots represent the average of the first 3 CSs of every day. Results are expressed in % of time during CS presentations, except in rearing behavior where the first plot expresses the average number of rearing bouts during CS presentations. **a.** Across extinction training, Yoked rats present significantly higher rearing counts (Q(1, 14) = 10.15, p = 0.006, η_p_^2^ = 0.219) and rearing duration (Q(1, 9.94) = 10.223, p = 0.01, η_p_^2^ = 0.333) compared to Avoiders in Experiment 1. **b.** Both groups significantly reduce their rearing duration (Q(3, 9.88) = 3.94, p = 0.043, η_p_^2^ = 0.276) across extinction training in Experiment 2. **c.** Rearing events and duration in Experiment 3 are not significantly different across group and training session. *p < 0.05, ** p < 0.01.

Fig S4. Extinction with platform in Experiment 1

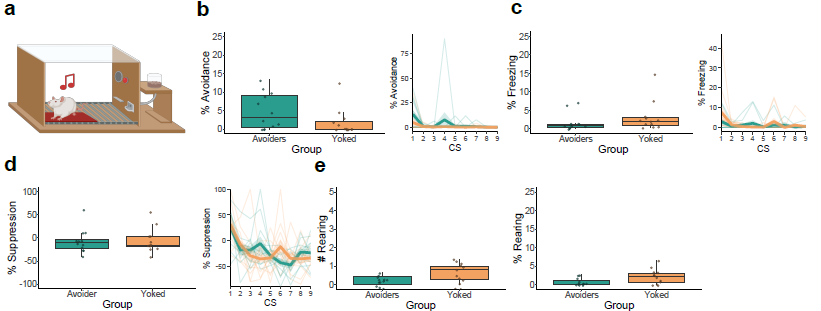

*Fig S4.* Extinction with platform results for Experiment 1. The box plots represent the average of the first 3 CSs. The bold lines in the trial-by-trial plots represent the mean and the surrounding shaded area the standard error of the mean. Results are expressed in % of time during CS presentations, except in rearing behavior where the first plot expresses the average number of rearing bouts during CS presentations. **a.** Graphical representation of the extinction with platform session. **b.** Both groups show similar low levels of avoidance in the extinction session with platform. **c.** An interaction between group and block (Q(2, 8.88) = 6.68, p = 0.017, η_p_^2^ = 0.046) in freezing in the extinction session with platform. Further analyses were not significant after multiple-testing correction. **d.** Both groups similarly reduced their suppression of lever pressing over the three blocks (Q(2, 12.11) = 6.89, p = 0.01, : η_p_^2^ = 0.331). **e.** Both groups similarly reduced their amount (Q(2, 11.93) = 7.77, p = 0.007, η_p_^2^ = 0.279) and duration of rearing (Q(2, 11.72) = 6.52, p = 0.012, η_p_^2^ = 0.215) over the blocks.

Fig S5. Rearing in reinstatement test Experiment 2 and 3

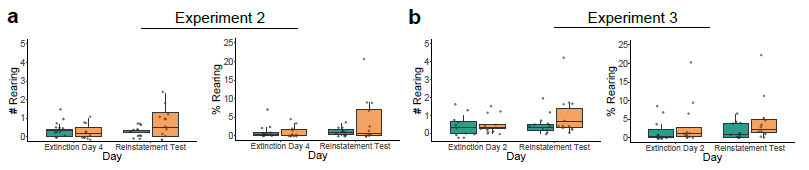

*Fig S5.* Rearing events and duration in reinstatement test Experiments 2 and 3. The box plots represent the average of the first 3 CSs of the reinstatement test compared to the preceding extinction day. Results are expressed in % of time during CS presentations, except in rearing behavior where the first plot expresses the average number of rearing bouts during CS presentations. **a.** Both groups showed similar rearing behavior in Experiment 2. **b** Both groups showed similar rearing behavior in Experiment 3.

Fig S6. Platform switch Experiment 2 and 3

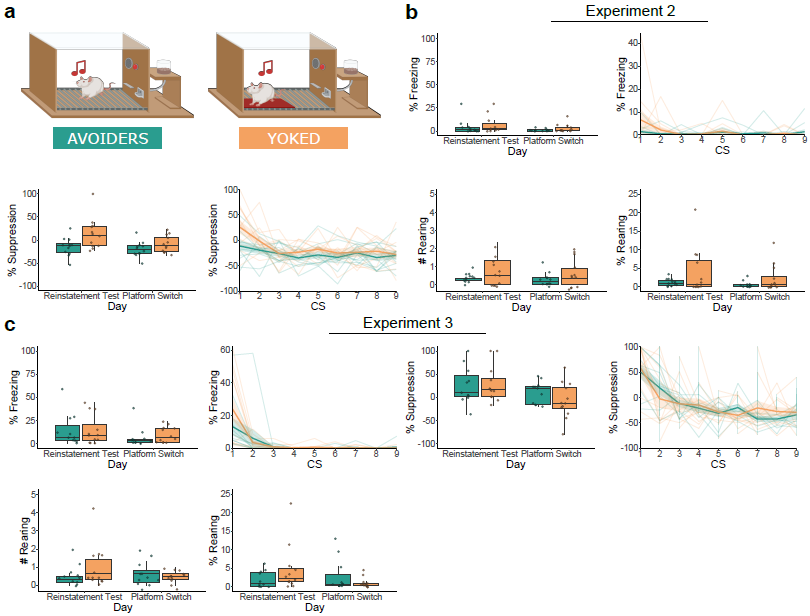

*Fig S6.* Platform switch results for Experiment 2 and 3. The box plots represent the average of the first 3 CSs of the platform switch session compared to the preceding reinstatement test. The bold lines in the trial-by-trial plots represent the mean and the surrounding shaded area the standard error of the mean. Results are expressed in % of time during CS presentations, except in rearing behavior where the first plot expresses the average number of rearing bouts during CS presentations. **a.** Graphical representation of the platform switch session. **b.** Avoider and Yoked rats show similar freezing, suppression and rearing behavior in the platform switch session in Experiment 2. **c.** Both groups show similar freezing, suppression and rearing behavior in the platform switch session in Experiment 3.

Fig S7. Alternative floor session Experiment 2

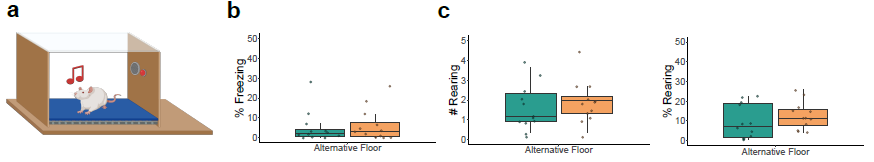

*Fig S7.* Alternative floor results for Experiment 2. The box plots represent the average of the first 3 CSs. Results are expressed in % of time during CS presentations, except in rearing behavior where the first plot expresses the average number of rearing bouts during CS presentations. **a.** Graphical representation of the alternative floor session. **b.** Both groups show similar freezing in the alternative floor session in Experiment 2. **c.** Both groups showed similar amount and duration of rearing in the alternative floor session in Experiment 2.

Fig S8. Rearing in spontaneous recovery test Experiment 3

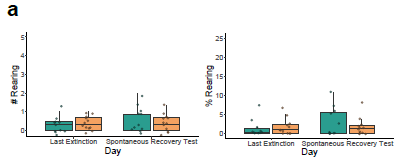

*Fig S8.* The box plots represent the average of the first 3 CSs of the spontaneous recovery test compared to the preceding extinction session. The first plot expresses the number of rearing bouts during those CS presentations and in the second plot results are expressed in % of time during CS presentations. Both groups showed similar amount and duration of rearing during the spontaneous recovery test in Experiment 3.

**Supplementary tables**

Table S1. Pavlovian training.

| Experiment | Measure | Statistical Test | Result | Effect size |
| --- | --- | --- | --- | --- |
| 3 | Freezing | T-test | t(20.6) = 0.298, p = 0.768 | d = 0.124 |
|  | Suppression of lever pressing | T-test | t(19.8) = 2.64, **p = 0.016** | d = 1.1 |
|  | Rearing events | Mann Whitney U test | W = 50, p = 0.331 | r = 0.209 |
|  | Rearing duration | T-test | t(16.7) = 1.02, p = 0.324 | d = 0.428 |

Table S2. Additional behaviors during avoidance training.

| Experiment | Measure | Statistical Test | Result | Effect size |
| --- | --- | --- | --- | --- |
| 1 | Suppression of lever pressing | Mixed ANOVA | Group: F(1, 22) = 0, p = 0.954  Day: F(1, 22) = 18.32, **p < 0.001**  G*D: F(1, 22) = 1.65, p = 0.213 | Group: η_p_^2^ < 0.001  Day: η_p_^2^ = 0.454  G*D: η_p_^2^ = 0.07 |
|  | Rearing events | Non-parametric mixed ANOVA | Group: Q(1, 13.43) = 4.95, **p = 0.044**  Day: Q(1, 13.05) = 18.32, **p < 0.001**  G*D: Q(1, 13.05) = 1.65, p = 0.213 | Group: η_p_^2^ = 0.267  Day: η_p_^2^ = 0.281  G*D: η_p_^2^ = 0.128 |
|  | Rearing duration | Non-parametric mixed ANOVA | Group: Q(1, 9.56) = 6.96, **p = 0.026**  Day: Q(1, 9.35) = 0.55, p = 0.477  G*D: Q(1, 9.35) = 0.03, p = 0.868 | Group: η_p_^2^ = 0.234  Day: η_p_^2^ < 0.001  G*D: η_p_^2^ < 0.001 |
| 2 | Suppression of lever pressing | Non-parametric mixed ANOVA | Group: Q(1, 14) = 0.108, p = 0.747  Day: Q(1, 12.58) = 161.70, **p < 0.001**  G*D: Q(1, 12.58) = 0.40, p = 0.538 | Group: η_p_^2^ = 0.019  Day: η_p_^2^ = 0.854  G*D: η_p_^2^ = 0.045 |
|  | Rearing events | Mixed ANOVA | Group: F(1, 22) = 3.65, p = 0.069  Day: F(1, 22) = 2.08, p = 0.163  G*D: F(1, 22) = 0.75, p = 0.396 | Group: η_p_^2^ = 0.142  Day: η_p_^2^ = 0.086  G*D: η_p_^2^ = 0.033 |
|  | Rearing duration | Non-parametric mixed ANOVA | Group: Q(1, 12.45) = 1.45, p = 0.251  Day: Q(1, 13.99) = 0.61, p = 0.447  G*D: Q(1, 13.99) = 0.09, p = 0.773 | Group: η_p_^2^ = 0.086  Day: η_p_^2^ = 0.043  G*D: η_p_^2^ < 0.001 |
| 3 | Suppression of lever pressing | Non-parametric mixed ANOVA | Group: Q(1, 6.33) = 14.59, **p = 0.008**  Day: Q(1, 6.33) = 5.75, p = 0.051  G*D: Q(1, 6.33) = 4.33, p = 0.08 | Group: η_p_^2^ = 0.38  Day: η_p_^2^ = 0.262  G*D: η_p_^2^ = 0.073 |
|  | Rearing events | Mixed ANOVA | Group: F(1, 21) = 4.68, **p = 0.042**  Day: F(1, 21) = 6.66, **p = 0.017**  G*D: F(1, 21) = 0, p = 0.984 | Group: η_p_^2^ = 0.182  Day: η_p_^2^ = 0.241  G*D: η_p_^2^ < 0.001 |
|  | Rearing duration | Non-parametric mixed ANOVA | Group: Q(1, 8.76) = 4.49, p = 0.064  Day: Q(1, 10.03) = 6.03, **p = 0.034**  G*D: Q(1, 10.03) = 0, p = 0.981 | Group: η_p_^2^ = 0.163  Day: η_p_^2^ = 0.105  G*D: η_p_^2^ = 0.01 |

Table S3. Rearing during extinction training

| Experiment | Measure | Statistical Test | Result | Effect size |
| --- | --- | --- | --- | --- |
| 1 | Rearing events | Non-parametric mixed ANOVA | Group: Q(1, 14) = 10.15, **p = 0.006**  Day: Q(3, 10.42) = 0.283, p = 0.8363  G*D: Q(3, 10.42) = 1.77, p = 0.213 | Group: η_p_^2^ = 0.219  Day: η_p_^2^ = 0.014  G*D: η_p_^2^ = 0.061 |
|  | Rearing duration | Non-parametric mixed ANOVA | Group: Q(1, 9.94) = 10.223, **p = 0.01**  Day: Q(3, 7.74) = 2.13, p = 0.176  G*D: Q(3, 7.74) = 2.09, p = 0.182 | Group: η_p_^2^ = 0.333  Day: η_p_^2^ = 0.135  G*D: η_p_^2^ = 0.113 |
| 2 | Rearing events | Non-parametric mixed ANOVA | Group: Q(1, 13.65) = 1.84, p = 0.197  Day: Q(3, 10.88) = 2.28, p = 0.137  G*D: Q(3, 10.88) = 0.24, p = 0.869 | Group: η_p_^2^ = 0.008  Day: η_p_^2^ = 0.225  G*D: η_p_^2^ = 0.024 |
|  | Rearing duration | Non-parametric mixed ANOVA | Group: Q(1, 13.68) = 1.52, p = 0.239  Day: Q(3, 9.88) = 3.94, **p = 0.043**  G*D: Q(3, 9.88) = 0.45, p = 0.725 | Group: η_p_^2^ < 0.001  Day: η_p_^2^ = 0.276  G*D: η_p_^2^ = 0.008 |
|  | Rearing duration | Pairwise comparisons with Wilcoxon rank sum test with Bonferroni correction | Day 1-2: **p = 0.002**  Day 1-3: **p = 0.002**  Day 2-3: p = 1  Day 1-4: **p = 0.036**  Day 2-4: p = 1  Day 3-4: p = 1 |  |
| 3 | Rearing events | Non-parametric mixed ANOVA | Group: Q(1, 9.30) = 1.07, p = 0.327  Day: Q(1, 12) = 2.64, p = 0.130  G*D: Q(1, 12) = 1.58, p = 0.232 | Group: η_p_^2^ = 0.034  Day: η_p_^2^ = 0.122  G*D: η_p_^2^ = 0.062 |
|  | Rearing duration | Non-parametric mixed ANOVA | Group: Q(1, 11.8) = 0.83, p = 0.382  Day: Q(1, 11.08) = 3.81, p = 0.077  G*D: Q(1, 11.08) = 0.19, p = 0.667 | Group: η_p_^2^ = 0.028  Day: η_p_^2^ = 0.059  G*D: η_p_^2^ = 0.007 |

Table S4. Extinction with platform in Experiment 1

| Experiment | Measure | Statistical Test | Result | Effect size |
| --- | --- | --- | --- | --- |
| 1 | Avoidance | Non-parametric mixed ANOVA | Group: Q(1, 7.69) = 2.36, p = 0.164  Block: Q(2, 7.18) = 3.59, p = 0.083  G*B: Q(2, 7.18) = 1.68, p = 0.252 | Group: η_p_^2^ = 0.116  Block: η_p_^2^ = 0.143  G*B: η_p_^2^ = 0.04 |
|  | Freezing | Non-parametric mixed ANOVA | Group: Q(1, 13) = 2.7, p = 0.124  Block: Q(2, 8.88) = 10.96, **p = 0.004**  G*B: Q(2, 8.88) = 6.68, **p = 0.017** | Group: η_p_^2^ = 0.021  Block: η_p_^2^ = 0.169  G*B: η_p_^2^ = 0.046 |
|  | Freezing | Simple effects by day: Wilcoxon signed ranks test | Block 1 – Group: V= 39.5, p = 0.064  Block 2 – Group: V= 60.5, p = 0.497  Block 3 – Group: V= 92.5, p = 0.217 | Block 1 – Group: r = 0.384  Block 2 – Group: r = 0.145  Block 3 – Group: r = 0.258 |
|  | Suppression of lever pressing | Non-parametric mixed ANOVA | Group: Q(1, 13.2) = 0.19, p = 0.664  Block: Q(2, 12.11) = 6.89, **p = 0.01**  G*B: Q(2, 12.11) = 0.19, p = 0.829 | Group: η_p_^2^ = 0.003  Block: η_p_^2^ = 0.331  G*B: η_p_^2^ = 0.003 |
|  | Suppression of lever pressing | Pairwise comparisons with Wilcoxon rank sum test with Bonferroni correction | Block 1-2: **p = 0.003**  Block 1-3: **p < 0.001**  Block 2-3: p = 1 |  |
|  | Rearing events | Non-parametric mixed ANOVA | Group: Q(1, 13.16) = 3.06, p = 0.103  Block: Q(2, 11.93) = 7.77, **p = 0.007**  G*B: Q(2, 11.93) = 2.76, p = 0.103 | Group: η_p_^2^ = 0.144  Block: η_p_^2^ = 0.279  G*B: η_p_^2^ = 0.185 |
|  | Rearing events | Pairwise comparisons with Wilcoxon rank sum test with Bonferroni correction | Block 1-2: **p = 0.005**  Block 1-3: p = 0.335  Block 2-3: p = 0.11 |  |
|  | Rearing duration | Non-parametric mixed ANOVA | Group: Q(1, 12.17) = 3.1, p = 0.103  Block: Q(2, 11.72) = 6.52, **p = 0.012**  G*B: Q(2, 11.72) = 2.53, p = 0.122 | Group: η_p_^2^ = 0.082  Block: η_p_^2^ = 0.215  G*B: η_p_^2^ = 0.147 |
|  | Rearing duration | Pairwise comparisons with Wilcoxon rank sum test with Bonferroni correction | Block 1-2: p = 0.006  Block 1-3: p = 0.579  Block 2-3: p = 0.098 |  |

Table S5. Rearing during the reinstatement test

| Experiment | Measure | Statistical Test | Result | Effect size |
| --- | --- | --- | --- | --- |
| 2 | Rearing events | Non-parametric mixed ANOVA | Group: Q(1, 8.89) = 0.36, p = 0.565  Day: Q(1, 10.84) = 1.68, p = 0.222  G*D: Q(1, 10.84) = 1.68, p = 0.222 | Group: η_p_^2^ = 0.055  Day: η_p_^2^ = 0.098  G*D: η_p_^2^ = 0.162 |
|  | Rearing duration | Non-parametric mixed ANOVA | Group: Q(1, 8.23) = 0.48, p = 0.508  Day: Q(1, 8.32) = 2.02, p = 0.19  G*D: Q(1, 8.32) = 0.73, p = 0.418 | Group: η_p_^2^ = 0.081  Day: η_p_^2^ = 0.105  G*D: η_p_^2^ = 0.105 |
| 3 | Rearing events | Non-parametric mixed ANOVA | Group: Q(1, 9.06) = 1.23, p = 0.296  Day: Q(1, 11.67) = 3.05, p = 0.107  G*D: Q(1, 11.67) = 1.83, p = 0.201 | Group: η_p_^2^ = 0.051  Day: η_p_^2^ = 0.120  G*D: η_p_^2^ = 0.079 |
|  | Rearing duration | Non-parametric mixed ANOVA | Group: Q(1, 11.07) = 1.04, p = 0.329  Day: Q(1, 11.97) = 1.44, p = 0.253  G*D: Q(1, 11.97) = 0.05, p = 0.823 | Group: η_p_^2^ = 0.098  Day: η_p_^2^ = 0.009  G*D: η_p_^2^ = 0.006 |

Table S6. Platform switch session

| Experiment | Measure | Statistical Test | Result | Effect size |
| --- | --- | --- | --- | --- |
| 2 | Freezing | Mann Whitney U test | W = 45.5, p = 0.119 | r = 0.324 |
|  | Suppression of lever pressing | T-test | t(21.8) = -1.63, p = 0.117 | d = -0.67 |
|  | Rearing events | Mann Whitney U test | W = 56, p = 0.346 | r = 0.199 |
|  | Rearing duration | Mann Whitney U test | W = 52.5 p = 0.255 | r = 0.239 |
| 3 | Freezing | Mann Whitney U test | W = 50, p = 0.34 | r = 0.205 |
|  | Suppression of lever pressing | T-test | t(18.2) = 1.48, p = 0.157 | d = 0.603 |
|  | Rearing events | Mann Whitney U test | W = 72, p = 0.726 | r = 0.079 |
|  | Rearing duration | Mann Whitney U test | W = 72, p = 0.733 | r = 0.077 |

Table S7. Alternative floor session

| Experiment | Measure | Statistical Test | Result | Effect size |
| --- | --- | --- | --- | --- |
| 2 | Freezing | Mann Whitney U test | W = 65, p = 0.706 | r = 0.083 |
|  | Rearing events | T-test | t(22) = -0.810, p = 0.426 | d = -0.173 |
|  | Rearing duration | Mann Whitney U test | W = 50, p = 0.214 | r = 0.259 |

Table S8. Rearing during the spontaneous recovery test

| Experiment | Measure | Statistical Test | Result | Effect size |
| --- | --- | --- | --- | --- |
| 3 | Rearing events | Non-parametric mixed ANOVA | Group: Q(1, 10.43) = 0.07, p = 0.796  Day: Q(1, 10.41) = 0.12, p = 0.74  G*D: Q(1, 10.41) = 0.12, p = 0.74 | Group: η_p_^2^ < 0.001  Day: η_p_^2^ = 0.057  G*D: η_p_^2^ = 0.034 |
|  | Rearing duration | Non-parametric mixed ANOVA | Group: Q(1, 8.10) = 0.03, p = 0.866  Day: Q(1, 8.67) = 0.88, p = 0.373  G*D: Q(1, 8.67) = 0.92, p = 0.364 | Group: η_p_^2^ = 0.009  Day: η_p_^2^ = 0.049  G*D: η_p_^2^ = 0.01 |
